# Septin-microtubule association requires a MAP-like motif unique to Sept9 isoform 1 embedded into septin octamers

**DOI:** 10.1101/2021.04.06.438596

**Authors:** Mira Kuzmić, Gerard Castro Linares, Jindřiška Leischner Fialová, François Iv, Danièle Salaün, Alex Llewellyn, Maxime Gomes, Mayssa Belhabib, Yuxiang Liu, Keisuke Asano, Taro Tachibana, Gijsje Koenderink, Ali Badache, Manos Mavrakis, Pascal Verdier-Pinard

**Author notes:** current affiliation: Department of Biology, University of Copenhagen, Copenhagen, Denmark (Jindřiška Leischner Fialová).

## Abstract

Septins, a family of GTP-binding proteins assembling into higher order structures, interface with the membrane, actin filaments and microtubules, which positions them as important regulators of cytoarchitecture. Septin 9 (Sept9), which is frequently overexpressed in tumors and mutated in hereditary neuralgic amyotrophy (HNA), mediates the binding of septins to microtubules, but the molecular determinants of this interaction remained uncertain. We demonstrate that a short MAP-like motif unique to Sept9 isoform 1 (Sept9_i1) drives septin octamer-microtubule interaction in cells and *in vitro* reconstitutions. Septin-microtubule association requires polymerizable septin octamers harboring Sept9_i1. Although outside of the MAP-like motif, HNA mutations abrogates this association, identifying a putative regulatory domain. Removal of this domain from Sept9_i1 sequesters septins on microtubules, promotes microtubule stability and alters actomyosin fiber distribution and tension. Thus, we identify key molecular determinants and potential regulatory roles of septin-microtubule interaction, paving the way to deciphering the mechanisms underlying septin associated pathologies.

## Introduction

Microtubules and actin filaments are cytoskeletal polymers involved in key cellular functions. Microtubules assemble from tubulin dimers to form hollow tubes that have the ability to switch between growing and shortening phases, a process called dynamic instability. Microtubule diversity arises, from tubulin isotypes, from post-translational modifications, including acetylation and glutamylation, often associated with stable sub-populations of microtubules, and from associations with specific microtubule-associated proteins (MAPs) that modulate microtubule dynamics^1^. Microtubules are critically involved in the segregation of chromosomes during cell division and the directed transport of various intracellular cargoes^2^. Actin monomers assemble into double helical strands that in cells are often cross-linked into bundles and networks^3^. In particular, the presence of associated myosins confers contractility to these actin fiber assemblies, that is essential during both cell division and migration. Such bundled actomyosin fibers form stress fibers that are qualified as peripheral and ventral when running near the cortex and the cell base, respectively. Stress fibers connect focal adhesions and the perinuclear actin cap linked to the top of the nucleus controlling its position and shape^4^. Several biological processes depend on the coordinated regulation of the actin and microtubule cytoskeletons, mediated by a diversity of molecular crosstalk between the two cytoskeletons. Direct and indirect contacts are involved, for instances, in cytokinetic furrow positioning, cell migration steering or maturation of neuronal dendritic spines^5^.

Septins are GTP-binding protein that form heteropolymeric complexes associating with membranes, actin filaments and a subset of microtubules^6^. Like yeast and Drosophila septins, human septin hexameric and octameric protomers can polymerize into filaments^7–13^. Septins are thought to function as diffusion barriers for protein compartmentalization or as scaffolds for protein-protein interactions during cells division and in an increasing number of processes during interphase^14–16^. During cytokinesis, septins are recruited to the sub-membrane cortex by anillin^17,18^ and co-localize with the constricting actomyosin ring, prior to the specification and the maintenance of the ingression furrow position by microtubules^19,20^. Septins appear also to control abscission^17,21–25^ that achieves cell division. It is still unclear how human septins interface with microtubules and the acto-myosin networks. Septins form a collar structure at the neck of dendritic spines, nanotubule-like protrusions and micro-tentacles to guide microtubules^26–29^. In adherent cells, septins concentrate on ventral stress fibers, and occasionally on perinuclear microtubules, depending on the cell type. Despite studies showing that septins regulate the traffic of motor proteins, like kinesin, the binding of MAPs and the guiding of microtubules in polarizing epithelia^30^, the functional consequences of septin-microtubule interaction are still largely unknown.

The 13 human septin genes encode four protein sequence homology-groups named after Sept2, Sept6, Sept7 and Sept3; septins from each group interact with each other in a specific order to form hexamers and octamers^22,31,32^. Septin octamers differ from hexamers by the addition of septins from the Sept3 group, with their arrangement recently determined as Sept2-Sept6-Sept7-Sept3-Sept3-Sept7-Sept6-Sept2^10,33^. Septins include a conserved GTP-binding domain (G domain) flanked by variable N- and C-termini. Septins assemble by the G domain on one side forming the G:G interface and by the N- and C- termini forming the NC:NC interface on the opposite side. This results in (_NC_Sept2_G:G_Sept6_NC:NC_Sept7_G:G_Sept7_NC:NC_Sept6_G:G_Sept2_NC_) hexamers and (_NC_Sept2_G:G_Sept6_NC:NC_Sept7_G:G_Sept3_NC:NC_Sept3_G:G_Sept7_NC:NC_Sept6_G:G_Sept2_NC_) octamers that can polymerize via a universal Sept2_NC_:_NC_Sept2 interface, that is labile in presence of high salt concentrations. In the Sept3 group, Sept9 has been the most studied because it is ubiquitously expressed, and often overexpressed in tumors^34^. Interestingly Sept9 is mutated in a large number of hereditary neuralgic amyotrophy (HNA) patients^35–37^. Alternative splicing resulting in alternative translation start sites gives rise to five Sept9 isoforms (Sept9_i1 to_i5) differing by the length and/or sequence of their N-terminus and by their functions^38–40^. Sept9_i1, _i2 and _i3 long isoforms share a common structurally disordered N-terminal domain (common N-ter) that was proposed, based on *in vitro* studies, to mediate binding to microtubules^41^. However, several studies indicate that only the isoform 1 of Sept9 (Sept9_i1) drives the association of septins with microtubules^24,32,40,42^. In order to understand the contribution of septin-microtubule interaction to cell physiology and disease, it is critical to identify and characterize precisely the molecular determinants of this interaction.

In the present study, we show that septin association with microtubules requires a MAP-like motif specific of Sept9_i1 and the integration of Sept9_i1 within polymerized septin octamers. *In vitro* reconstitution allowed us to demonstrate a direct and specific interaction of Sept9_i1-harboring septin octamers with microtubules, which slows down depolymerization of microtubules. Based on this molecular characterization, we designed specific genetic variants, including HNA-like mutations, which modulate Sept9_i1-microtubule interaction and we show that relocalizing septins to microtubules disturbs stress fiber distribution and tension.

## Results

### Different Sept9 isoform expression profiles are associated with distinct octamers/hexamers ratios and cytoskeletal localization

We have analyzed septin expression profiles in commonly used cell models, namely U2OS osteosarcoma, HeLa cervical carcinoma cells, RPE1 retinal epithelial cells and SKBr3 breast carcinoma cells (Fig. 1a). Sept7 being unique in its group, it is essential for the formation of hexamers and octamers (Supplementary Fig. 1d). The four cell lines examined expressed very similar levels of Sept7. In the Sept2 group, Sept2 was expressed in all cell lines and Sept5 was only detected in U2OS cells. Expression of septins from the Sept6 group was more variable, except for Sept8 that was expressed at similar levels in all cell lines. U2OS and HeLa cells expressed similar levels of total Sept9. However, U2OS cells expressed mostly isoform 3, whereas HeLa cells expressed mostly isoform 1. RPE1 and SKBr3 cells expressed higher levels of Sept9 with similar levels of Sept9_i1 and Sept9_i2, and no or very low levels of Sept9_i3.

**Fig 1.**
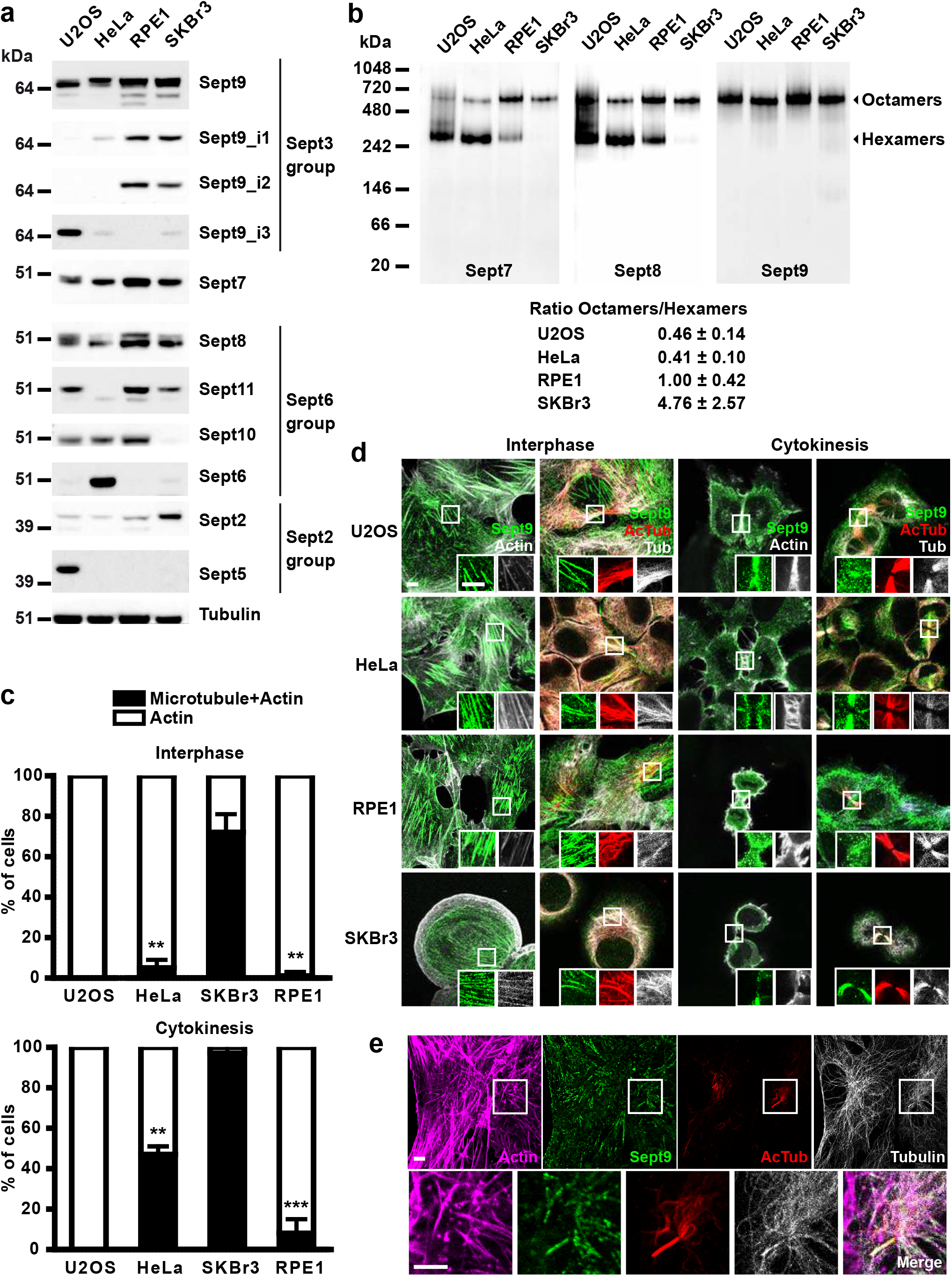
Sept9 expression, incorporation in octamers and localization in human cell lines. **a** Western blots of total protein extracts from U2OS, Hela, RPE1 and SKBr3 cells probed with antibodies against septins and tubulin. **b** Native extracts from the same cell lines were analyzed by Western blotting for their content in both septin hexamers and octamers using antibodies against constitutively required Sept7 and ubiquitously expressed Sept8; septin octamers were also specifically labeled with a pan Sept9 antibody. Relative content in septin octamers and hexamers in each cell line based on three independent Sept7 WB is presented below the blots (mean ± SEM). **c** and **d** Cellular localization by immunocytochemistry of Sept9 in the same cell lines, during interphase and cytokinesis; in **c**, the percentages of cells displaying both co-localization of Sept9 with microtubules and co-localization with stress fibers (Microtubule + Actin) or only stress fibers (Actin) were determined from three independent experiments based on a total of 90 cells (30 cells per experiment). Unpaired, two-tailed t-test with Welch’s correction, ** p< 0.01, *** p<0.0005, Microtubule and Actin localization vs in SKBr3; in **d**, white scale bars correspond to 5 μm, insets are two-fold zoomed regions framed by a white square in the corresponding original images. **e** RPE1 cells were starved for 24 h and immunofluorescence was performed as in **c** and **d**; white scale bars correspond to 5 μm, a three-fold zoom of a region framed in white containing a primary cilium is presented below the original images.

To assess how Sept9 expression profiles affected its incorporation in septin hexamers and octamers, we extracted cells in conditions that preserved oligomers and performed native gel electrophoresis and analysis by Western blotting (Fig. 1b and Supplementary Fig. 1). Based on the detection of the essential Sept7 and pan-expressed Sept8, U2OS and HeLa cells contained more hexamers than octamers, as opposed to RPE1 and SKBr3 cells. This result is consistent with higher Sept9 expression levels in RPE1 and SKBr3 leading to higher levels of octamers. In addition, our results indicate that long Sept9 isoforms incorporate indifferently in octamers. We did not detect smaller septin assemblies containing Sept9 monomers or dimers, suggesting that, at endogenous levels, the entire pool of Sept9 is stably incorporated into octamers.

Next, we examined the localization of Sept9 in these cell lines either in interphase or in late cytokinesis (Fig. 1c, d). We observed that Sept9 was associated with actin fibers in all interphase cells, especially on ventral stress fibers (Fig. 1c, d). Interestingly, consistent with previous observation^40^, Sept9 co-localized with a subpopulation of acetylated microtubules in HeLa and SKBr3 cells; in cytokinetic cells, Sept9 concentrated cortically at the ingression furrow and on the thick bundles of acetylated microtubules contained in the intercellular bridge. In U2OS cells, which express primarily Sept9_i3, septins associated with the actin cytoskeleton but did not co-localize with microtubules. These results indicate that the association of septins with microtubules is dependent on the presence of octamers harboring Sept9_i1. It is less clear why in RPE1 cells, despite Sept9_i1 levels comparable to SKBr3 cells, there was no association of Sept9 with microtubules, except in primary cilia (Fig. 1e).

### Sept9_i1 interacts with microtubules via a specific MAP-like motif

The fact that only Sept9_i1, but not the closely related i2 and i3 isoforms, is required for septins to bind microtubules identifies the sequence formed by the first specific 25 amino acid residues of the Sept9_i1 protein, as specific to this isoform and as key for its interaction with microtubules^40^. This sequence being well conserved in vertebrates (Supplementary Fig. 2), it is also indicative of an important function. Searching for similar sequences in proteins related to the microtubule cytoskeleton by using the NCBI Blastp tool, we found the Arabidopsis thaliana AIR9 protein, a plant MAP involved in cytokinesis^43^. The sequence of AIR9 similar to the Sept9_i1 specific sequence was located in the AIR9-microtubule binding domain (AIR9 1-234)^43^. Alignment of these two sequences reveals a block of similar sequence that we named AIR-9-like (Fig. 2a). Another sequence, located upstream in the AIR9 N-terminus (aa 46-78), presents a lower similarity with the Sept9_i1 specific sequence (Fig. 2a). Strikingly, the AIR9-like block had similarities with canonical repeats found in the microtubule binding domains (MBDs) of the structural human MAPs, tau, MAP2 and MAP4, which are known to interact directly with the surface of microtubules (Fig. 2a). An upstream smaller block in the Sept9_i1 specific sequence showed similarities with the MAP4 R2 repeat. Thus, Sept9_i1 displays a putative MBD, similar to the MBD repeats of several MAPs and constituted by a MAP4 R2-like sequence followed by an AIR9-like sequence.

**Fig. 2.**
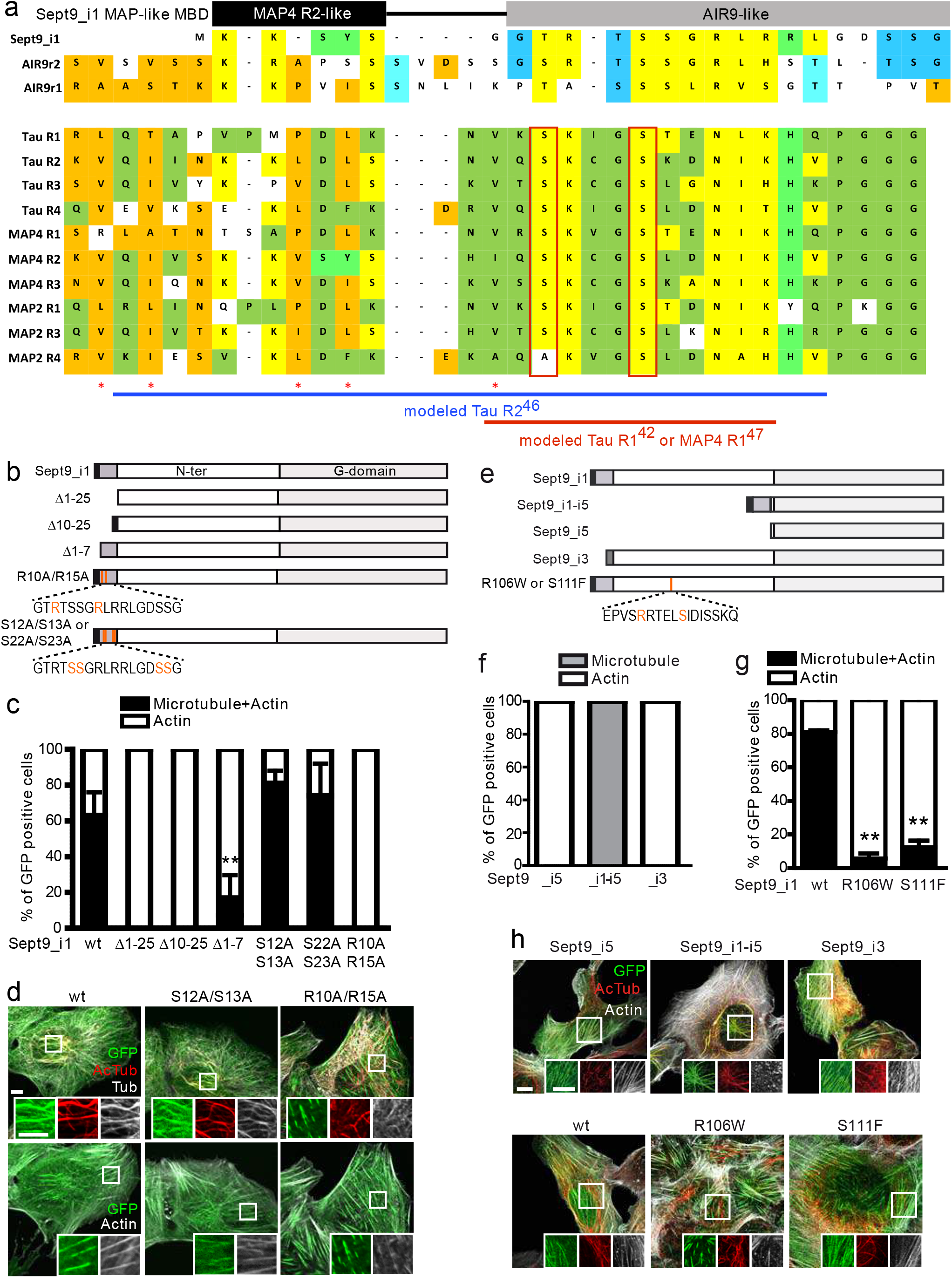
Identification of the microtubule binding domain in Sept9_i1. **a** Top part: human Sept9_i1 specific amino-acid N-terminal sequence (1-24) was aligned with the homologous plant *A. thaliana* AIR9 sequences (173-205 and 46-78) contained in the AIR-9 microtubule binding domain (MBD)^43^; bottom part: Sept9_i1 and AIR9 sequences in top part were aligned with sequences of tandem repeats involved in the binding of human structural MAPs (Tau, MAP4 and MAP2) to microtubules (ex: Tau 4R isoform positions of repeats are R1**_242-273_**, R2**_274-304_**, R3**_305-335_** and R4**_336-367_**). Alignments were adjusted by introducing gaps and similar aa residues were manually coded, orange: common to AIR9 and MAPs; yellow: common to Sept9_i1, AIR9 and MAPs; light green: common to Sept9_i1 and MAPs; dark green: common to MAPs; dark blue: common to Sept9_i1 and AIR9; light blue: common to AIR9 repeats. Important serine residues and hydrophobic residues of MAPs involved in MAP repeat-microtubule lattice interactions are framed in red and marked with a red asterisk, respectively. Horizontal blue line and red line indicate, respectively, the position of Tau R2 repeat and Tau R1/MAP4 R1 repeat modeled based on the cryo-EM structural determination of MAP repeated sequences bound on microtubules *in vitro*^46,47^. **b** Schematic representation of the Sept9_i1 mutants in the predicted MBD tagged with GFP at their C-terminus. **c**, **d** Co-localization of each Sept9 construct transfected in U2OS cells with microtubules or stress fibers during interphase was analyzed as described in Fig. 1 legend. Unpaired, two-tailed t-test with Welch’s correction, ** p< 0.01, Microtubule and Actin localization vs Sept9_i1 wt; white scale bars 5 μm, insets: three-fold zooms. **e** Schematic representation of Sept9 isoforms (Sept9_i1, _i5 and _i3), of the chimeric construct fusing the Sept9_i1 MBD to the N-terminus of Sept9_i5 (Sept9_i1-i5) and of Sept9_i1 HNA mutants, all tagged with GFP at their C-terminus. **f** Co-localization of Sept9_i5 or _i1-i5 or _i3, and **g** of Sept9 HNA point mutants with microtubules or stress fibers in U2OS transfected cells during interphase. Unpaired, two-tailed t-test with Welch’s correction, ** p< 0.01. **h** Images showing co-localization of Sept9 in U2OS cells transfected with constructs described in **e**; white scale bars 10 μm, insets: 1.4-fold zooms.

To unequivocally demonstrate that this motif is actually the MBD of Sept9_i1, we examined the localization of Sept9_i1 mutants (C-terminally tagged with GFP), presenting deletions or point mutations in the first 25 amino acid residues (Fig. 2b), in U2OS cells (Fig. 2c, d and Supplementary Fig. 3a-c). U2OS cells were chosen because they express very low levels of Sept9_i1 (Fig. 1a), and show no Sept9-microtubule co-localization (Fig. 1c, d). In addition, U2OS cells were amenable to simultaneous siRNA-mediated knockdown (KD) of Sept9 and expression of exogenous Sept9_i1-GFP constructs at near-endogenous Sept9 levels (Supplementary Fig. 3a). Wild type Sept9_i1 co-localized with actin in all cells and with microtubules in more than 60% of the cells. Sept9_i3 and the Δ1-25 Sept9_i1 mutant also associated with actin fibers, but never localized to microtubules (Fig. 2c, d, f and Supplementary Fig. 3b, d). This result is consistent with our previous observations of Sept9_i3 and Sept9_i1 Δ1-25 vs Sept9_i1 localization in SKBr3 cells (using N-terminally tagged constructs)^40^. Deletion of the AIR9-like portion (Δ10-25 mutant) or of the MAP4 R2-like portion (Δ1-7 mutant) totally or largely abrogated Sept9 binding to microtubules, respectively (Fig. 2c and Supplementary Fig. 3b). Next, we introduced point mutations in the AIR9-like portion (Fig. 2b). Independent mutations of two sets of contiguous serine residues (S12A/S13A and S22A/S23A), conserved in the AIR9 MBD, did not significantly alter the co-localization of Sept9 with microtubules (Fig. 2c, d and Supplementary Fig. 3b, c). Importantly, mutations of the two arginine residues (R10A/R15A), conserved in all vertebrates (Supplementary Fig. 2) and present in the AIR9 MBD, completely abrogated the binding of Sept9 to microtubules (Fig. 2c, d and Supplementary Fig. 3b, c). Collectively, our data show that the Sept9_i1 specific N-terminal sequence is an MBD analogous to the ones found in human MAPs and the plant AIR9 MAP. Of note, in contrast to MAP MBDs, the Sept9_i1 specific sequence is not tandem-repeated and lacks upstream conserved residues (Fig. 2a).

A previous report proposed, based on *in vitro* studies using recombinant polypeptides, that Sept9_i161-113 within the N-terminal region common to all Sept9 long isoforms (i1, i2 and i3) was the main contributor to microtubule binding^41^. To evaluate a potential contribution of Sept9 N-terminal region common to all long isoforms (common N-ter), we generated a chimeric Sept9 deleted of the entire common N-ter by fusing the isoform 1 specific sequence to Sept9_i5, the shortest Sept9 isoform, and named it Sept9_i1-i5 (Fig. 2e). Whereas Sept9_i5 localized strictly to actin fibers (Fig. 2f, h and Supplementary Fig. 3e), its fusion to the Sept9_i1 specific motif was sufficient to localize septins to microtubules in all interphase cells. Surprisingly, this was accompanied by the complete loss of Sept9 co-localization with actin fibers (Fig. 2f, h). Consistent with these findings, expression of Sept9_i1-i5 during cytokinesis increased strongly the percentage of cells showing Sept9 association with microtubule bundles at the intercellular bridge (Supplementary Fig. 3d).

Our results demonstrate that the Sept9_i1 MBD is required and sufficient to target Sept9 to microtubules, whereas the N-terminal region common to all long isoforms is dispensable. Interestingly the Sept9_i1-i5 is much more efficient than Sept9_1 for targeting septins to microtubules (Fig. 2f, h and Supplementary Fig. 3d), hinting that the common N-ter might have a negative regulatory function. Sept9 is mutated in hereditary neuralgic amyotrophy (HNA) patients. HNA-associated mutations are located in the Sept9 common N-ter^35^, but their impact on Sept9 function remains to be elucidated. We introduced separately the two most frequently described HNA mutations (R106W or S111F) in Sept9_i1 constructs and evaluated their impact on Sept9_i1 localization. Both Sept9_i1 R106W and S111F maintained co-localization with ventral stress fibers (Fig. 2g, h). However, both mutations suppressed the association with microtubules (Fig. 2g, h). This result confirms that the common N-ter, while not required for Sept9 microtubule binding per se, is an important regulatory domain and provides novel insights into the molecular etiology of HNA.

### Sept9_i1 must be included in a polymerized septin octamer to associate with microtubules

In contrast to typical MBDs in structural MAPs, the Sept9_i1 specific sequence does not have tandem repetitions. We hypothesized that the dimerization of Sept9_i1 in octamers via the NC interface brings two Sept9_i1 specific sequences in close proximity, to form a MBD mimicking tandem arrangements of repeats. To test this hypothesis and the oligomeric environment required for microtubule binding, we introduced point mutations into Sept9_i1 that should disturb the NC interface to generate tetramers, or the G interface to generate non-incorporated Sept9_i1^31^ (Fig. 3a). We then expressed these constructs in U2OS cells in place of endogenous Sept9 (KD via siRNA). Importantly, re-expression of wild type Sept9_i1-GFP restored close to endogenous levels of octamers and hexamers (Fig. 3a). As expected, most of the Sept9_i1 G interface mutant (Sept9_i1 G_mut_) did not incorporate in octamers (Fig. 3a). Instead, it migrated with an apparent molecular weight of 138 kDa, corresponding most likely to monomeric Sept9_i1, whose gel motility is retarded by the long and disordered N-terminus^32^ (Supplementary Fig. 1a). Based on previous studies^22,31,44^, we generated two Sept9_i1 NC interface mutants with mutations in the α0 helix either in the basic residue stretch (Sept9_i1 NC_mut1_) or upstream of this stretch in key hydrophobic residues (Sept9_i1 NC_mut2_). Sept9_i1 NC_mut1_ incorporated in tetramers and Sept9_i1 NC_mut2_ incorporated partly in tetramers, but mostly migrated like the monomeric protein (Fig. 3a), indicating that mutations in Sept9_i1 NC_mut2_ might affect both NC and G interfaces. Importantly, we observed no association of Sept9_i1 G or NC mutants with microtubules, but rather a diffuse cytosolic distribution (Fig. 3b). These results show that neither septin tetramers nor Sept9_i1 monomers bind to microtubules, and that the octameric septin context, which brings together two Sept9 units via their NC interface, is mandatory for Sept9_i1 association with microtubules.

**Fig. 3.**
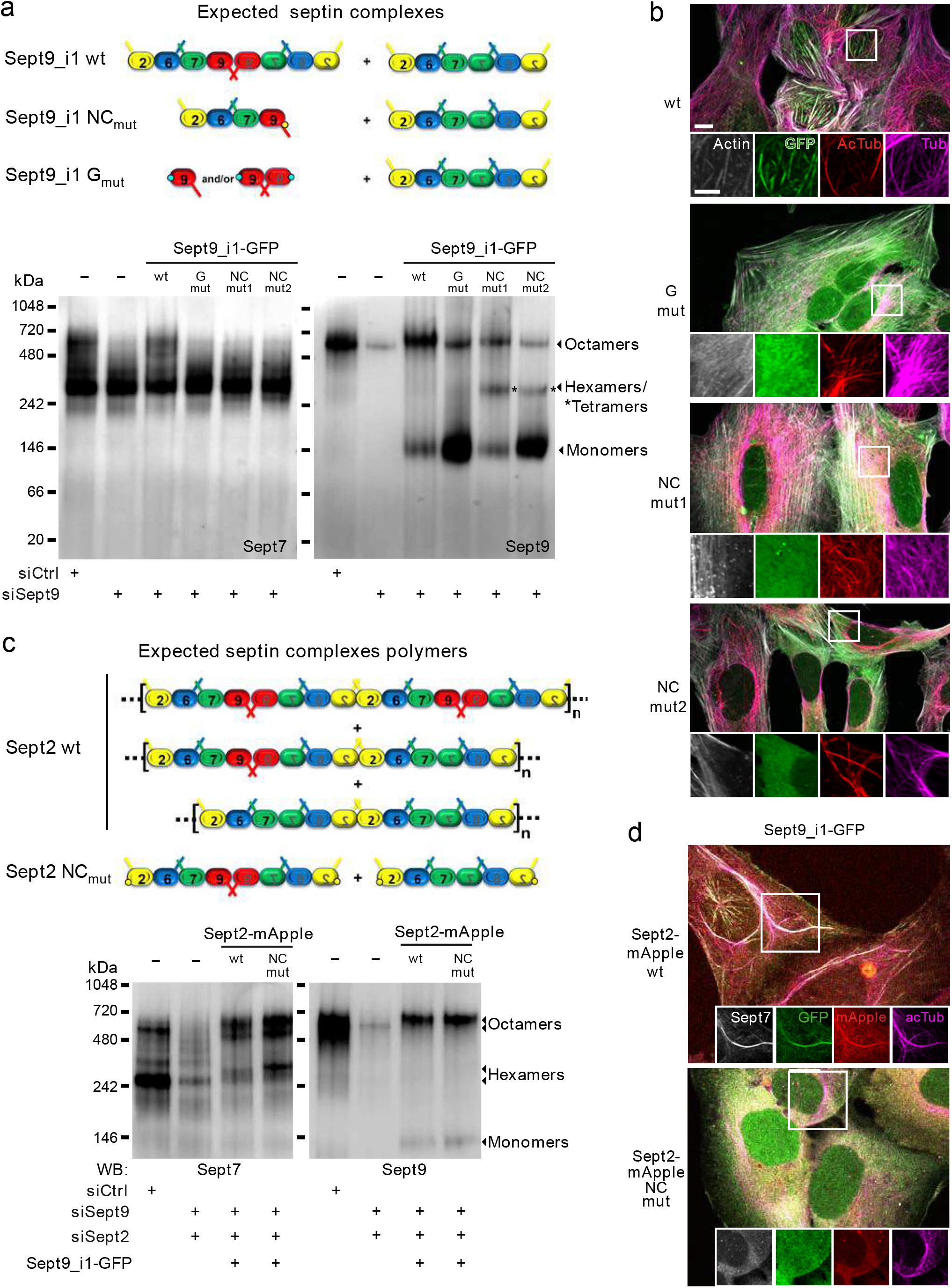
Dependence of septin association with microtubules on the incorporation of Sept9_i1 in polymerized septin octamers. **a** Top, predicted septin complexes when Sept9_i1 wt, or Sept9_i1 mutated on its G (G_mut_, blue dots) or NC interface (NC_mut_, yellow dot) is expressed in cells; bottom, Western blots of native gels resolving septin complexes present in U2OS cells expressing indicated siRNA and Sept9_i1 wt or Sept9_i1 interface mutants. **b** GFP-fluorescence based co-localization of Sept9_i1 wt or interface mutants with actin and microtubules; Sept9_i1 wt co-localized both with microtubule and actin fibers as expected, but non-octameric septin complexes containing an interface mutant Sept9_i1 did not. White scale bar 10 μm, insets: two-fold zooms of frames region. **c** Top, predicted septin complexes when Sept2 wt or Sept2 mutated on its NC interface (NC_mut_, yellow dots) is co-expressed with Sept9_i1 in U2OS cells; bottom, Western blots of native gels resolving septin complexes present in U2OS expressing indicated siRNAs and septin constructs. **d** mApple and GFP-based co-localization of Sept2 and Sept9_i1, respectively, with actin and microtubules; expression of Sept2 NC mutant prevented normal co-localization of octamers harboring Sept9_i1 with acetylated microtubules and actin fibers. Insets: individual fluorescence channel images of framed regions.

Next, we evaluated if septin octamers harboring Sept9_i1 bind microtubules by themselves or if they need to assemble into longer polymers via Sept2:Sept2 NC interfaces. Mutation of the Sept2 NC interface^31,45^ (Sept2 NC_mut_) should produce a mixture of isolated octamers and hexamers in cells (Fig. 3c). To evaluate its impact on Sept9_i1 localization, we knocked down both Sept9 and Sept2, and re-expressed Sept9_i1 (tagged with GFP to monitor Sept9 localization), together with wild type or Sept2 NC_mut_ (tagged with mApple to identify transfected cells). Using native gel separation, we verified that wild type Sept2 and Sept2 NC_mut_ generated the same relative expression levels of octamers and hexamers (Fig. 3c). However, when Sept2 NC_mut_ was expressed Sept9_i1 (and Sept2) no longer co-localized with microtubules (Fig. 3d). Thus, the association of Sept9_i1 with microtubules requires its incorporation in polymerizable octamers.

### Sept9_i1 associates with stable microtubules

Because of the sequence similarities between Sept9_i1 and MAP4 MBDs, the two proteins might use the same tubulin interdimer binding pocket at the surface of microtubules^46,47^. At endogenous Sept9_i1 levels in SKBr3 cells, we observed that MAP4 associated with the entire microtubule cytoskeleton, whereas Sept7 was concentrated on acetylated bundled microtubules (Fig. 4a). In U2OS cells transfected with Sept9_i1, the same differential distribution between MAP4 and Sept9_i1 was observed. When the Sept9_i1-i5 chimera, which showed a stronger affinity for microtubules, was expressed, MAP4 was absent on many microtubule bundles (Fig. 4b). These observations suggest that the binding sites of Sept9_i1 MBD and MAP repeats partially overlap on the microtubule lattice.

**Fig. 4.**
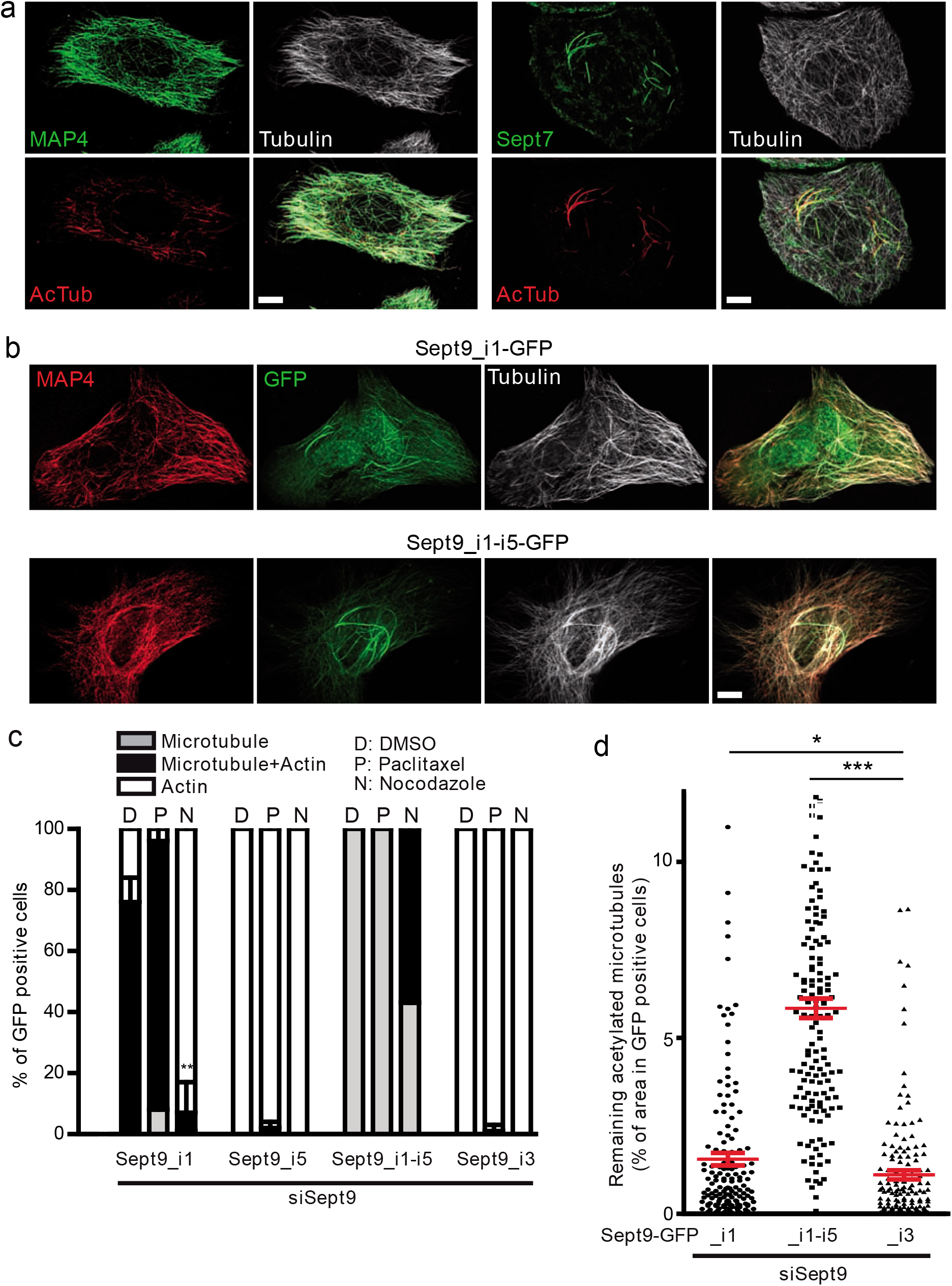
Sept9_i1 selective binding on, and stabilization of, microtubule bundles. **a** Differential co-localization of endogenous Sept7 vs MAP4 with microtubule cytoskeleton in SKBr3 cells. **b** Differential co-localization between endogenous MAP4 and transfected Sept9_i1-GFP or Sept9_i1-i5-GFP with microtubule cytoskeleton in U2OS cells. White scale bars 5 μm in merge images. **c** Analysis of the percent of U2OS cells KD for Sept9 and expressing the indicated Sept9-GFP constructs, displaying either microtubule or actin or mixed cytoskeleton co-localization with GFP during interphase, and treated either with DMSO vehicle alone (D) or 2 μM paclitaxel (P) or 10 μM nocodazole (N) for 2 hrs. Results are from the analysis on a total of 90 cells from three independent experiments based (30 cells per experiment). Unpaired, two-tailed t-test with Welch’s correction, ** p< 0.01, microtubule and actin localization vs DMSO. **d** Quantification of nocodazole-resistant acetylated microtubules (treatment as in **c**) in U2OS cells KD for Sept9 and expressing the indicated Sept9-GFP constructs. Results are from the analysis of a total of 150 cells from three independent experiments (50 cells per experiments). Unpaired, two-tailed t-test with Welch’s correction, * p<0.05, *** p<0.0005.

As we repeatedly observed the preference of Sept9_i1 for acetylated microtubules (Fig. 1d, e, Fig. 2d, h, Fig. 3b, d, Fig. 4a and Supplementary Fig. 3c, g), we wondered whether microtubule acetylation was a determinant of septin binding. We first examined whether the unexpected lack of Sept9_i1 association with microtubules in RPE1 cells (Fig. 1c, d) was associated with a deficit of microtubule acetylation. Tubulin acetylation levels in RPE1 cell line were indeed low, especially compared to those in another retina epithelial cell line (ARPE19), which expressed similar levels of Sept9_i1 (Fig. 5a) but showed frequent septin association with microtubules (Fig. 5b, supplementary Fig. 3f). Other posttranslational modifications (PTMs) of tubulins, such as polyglutamylation, that could affect Sept9_i1 binding^48,49^ were equally low in these cell lines (Fig. 5a). Moreover, treating RPE1 cells with paclitaxel, a drug that induces acetylation and bundling of microtubules in cells, induced the association of septins with microtubules (Fig. 5c). Of note, the paclitaxel-driven re-localization of septins to microtubules was dependent on the expression of Sept9_i1 (Fig. 4c and Fig. 5c). Thus, Sept9_i1 association with microtubules appears to be correlated with tubulin acetylation. To determine if paclitaxel-induced tubulin acetylation was responsible for re-localization of septins on microtubules, αTAT1, the major tubulin acetyl transferase^50^, was knocked down to prevent tubulin acetylation, prior to paclitaxel treatment (Fig. 5d). This experiment clearly showed that re-localization of septins on paclitaxel-bundled microtubules was independent of tubulin acetylation levels (Fig. 5d).

**Fig. 5.**
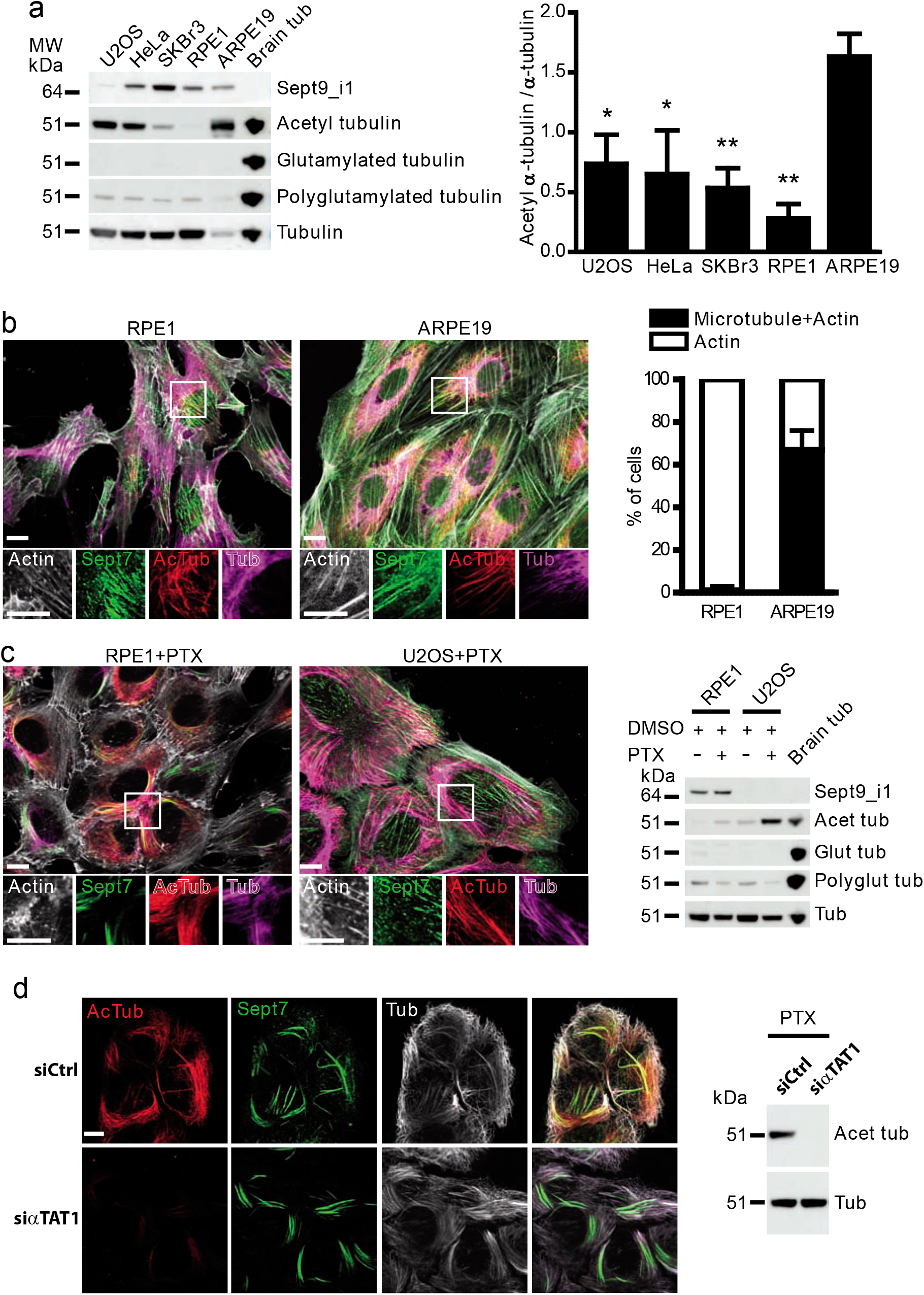
Dependence of Sept9_i1 binding to microtubules on their bundling but not on their acetylation or polyglutamylation. **a** Left: Western blots of total protein extracts from indicated cell lines, probed for Sept9_i1, α-tubulin acetylation on K40, glutamylation (one or two glutamate residues) and polyglutamylation of tubulin; as a positive control for these tubulin posttranslational modifications, an equivalent amount of purified porcine brain tubulin (Brain tub) was included in the last lane. Right: quantification of tubulin acetylation levels in cell lines based on acetylated α-tubulin and total α-tubulin Western blots. Results are from three independent determinations. Unpaired, two-tailed t-test with Welch’s correction, * p<0.05, ** p< 0.01, tubulin acetylation levels vs ARPE19. **b** Left: Co-localization by immunocytochemistry of Sept7 with microtubules and/or actin fibers in RPE1 and ARPE19 cells during interphase; white scale bars 10 μm, insets: two-fold zooms of framed regions. Right: quantification of this co-localization from three independent experiments on a total of 90 cells (30 cells per experiments). **c** Left: Co-localization by immunocytochemistry of Sept7 with microtubules bundles induced by paclitaxel treatment (PTX) and/or with actin fibers in RPE1 and ARPE19 cells; white scale bar 10 μm, insets: two-fold zooms of framed regions. Right: Western blots as described in **a** for total protein extracts of cells that were treated with vehicle alone (DMSO) or PTX. **d** Left: Co-localization by immunocytochemistry of Sept7 with microtubules bundles induced by paclitaxel treatment (PTX) in RPE1 cells either transfected with a control siRNA (siCtrl) or a siRNA against the α-tubulin acetyl transferase 1 (αTAT1); white scale bar 10 μm. Right: Western blot of total protein extracts from PTX treated RPE1 cells as described in the left part, showing down-regulation of α-tubulin acetylation upon αTAT1 KD.

Finally, we evaluated whether Sept9_i1, like MAPs, contributes to microtubule stabilization in the presence of nocodazole, a microtubule-depolymerizing agent. We observed that, whereas Sept9_i1 moderately stabilized microtubules, Sept9_i1-i5 strongly promoted the resistance of microtubules against nocodazole-induced depolymerization (Fig. 4c, d and Supplementary Fig. 3g). Of note, depolymerization of microtubules allowed Sept9_i1-i5 to re-localize to actin fibers in nearly 60% of the cells (Fig. 4c).

Collectively, these data establish that septin octamers harboring Sept9_i1 associate with a population of stable bundled microtubules in cells, independently of their acetylation levels, and contribute to microtubule stabilization.

### Recombinant septin octamers harboring Sept9_i1 interact directly and specifically with microtubules and slow down their depolymerization

In order to assess if recombinant octamers harboring (Oct_9i1) are able to bind directly to microtubules, we purified recombinant septin octamers (Sept2-Sept6-Sept7-Sept9_i1-Sept9_i1-Sept7-Sept6-Sept2 and Sept2-Sept6-Sept7-Sept9_i3-Sept9_i3-Sept7-Sept6-Sept2)^12^ (Supplementary Fig. 1b) and reconstituted dynamic microtubules made from purified tubulin α/β-heterodimers in the presence of each type of octamer. Total internal reflection fluorescence (TIRF) microscopy allowed visualizing both dynamic microtubules incorporating rhodamine-tagged tubulin and septin oligomers incorporating GFP-tagged Sept2 (Fig. 6a). Microtubules were polymerized from a 10 μM tubulin heterodimer solution and in the presence of 10 to 300 nM of octamers, i.e. at concentrations comparable to the tubulin α/β-heterodimer (~10 μM) and septin complex (~400 nM) concentrations measured in Hela cells^51^.

**Fig. 6.**
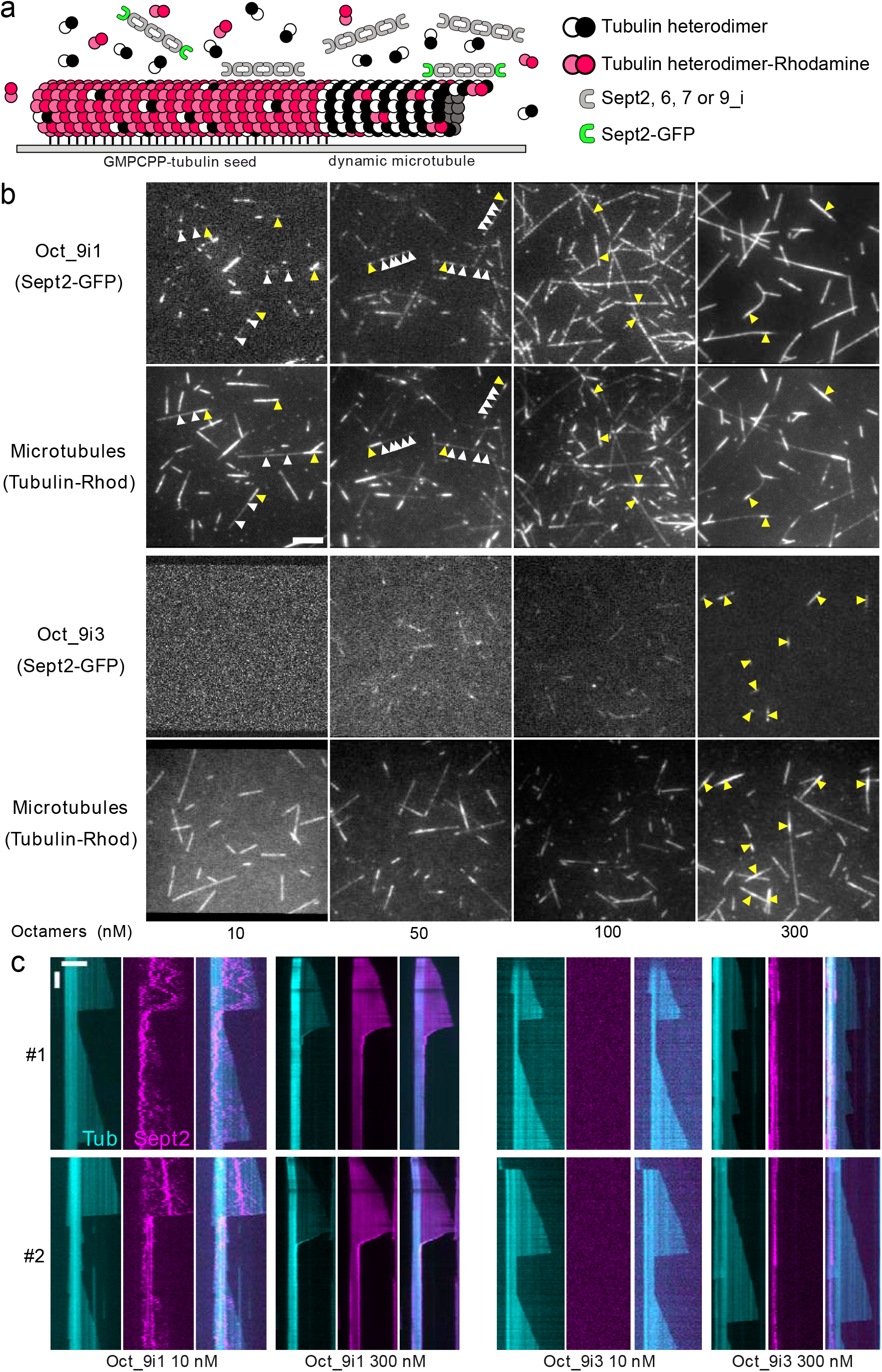
Recombinant septin octamers containing Sept9_i1 specifically bind microtubules *in vitro*. **a** Schematic representation of the *in vitro* assay. Biotinylated GMPCPP-stabilized microtubule seeds were immobilized on a streptavidin coated passivated glass coverslip, and free tubulin heterodimers (10 μM) along with septin octamers (nM concentrations as indicated containing 10% msfGFP-labeled septins) were added. The seeds were labelled with a higher percentage (30%) of rhodamine-tubulin heterodimers than the free tubulin added (3-6%) to be able to distinguish the seed from the dynamic microtubules while using the same fluorophore. **b** Septin octamers harboring either Sept9_i1 or _i3 subunits (Oct_9i1 or Oct_9i3) were introduced in the chamber at the indicated concentrations and their binding observed on dynamic microtubules (examples of sporadic binding indicated by white arrow heads) and stable microtubule seeds (examples indicated by yellow arrow heads). White scale bar 5 μm. **c** Examples (#1 and #2) of representative kymographs of microtubules observed as in **b** in the presence of the minimal (10 nM) or maximal (300 nM) septin octamer concentrations. False colors for rhodamine (Tub) and GFP (Sept2) fluorescence signals were used to optimize visually their merged image (third image from the left of each condition). Horizontal white scale bar 5 μm; vertical white scale bar 50 sec.

We clearly observed Oct_9i1 binding to microtubules at the lowest concentration (10 nM, Fig. 6b, c). Oct_9i1 diffused randomly on the lattice of dynamic microtubules (Fig. 6c and Supplementary Movie 1), and were also on the stable microtubule seeds. On kymographs, diffusion on microtubules during the growing phases was clearly apparent, with a reduced presence on the microtubule seeds (Fig. 6c). Of note, upon microtubule depolymerization, Oct_9i1 accumulated and remained on seeds, then diffused again on the microtubule upon its regrowth (Fig. 6c). At 10 nM concentration, there were no detectable recombinant octamers harboring Sept9_i3 (Oct_9i3) on dynamic or stable seed microtubules. Decoration of microtubules increased with increasing concentrations of Oct_9i1, until saturation at 300 nM (Fig. 6b, c and Supplementary Movie 1). In comparison, Oct_9i3 only showed significant binding at the highest concentration (300 nM), which was restricted to the stable microtubule seeds (Fig. 6b, c and Supplementary Movie 2). Altogether these results indicate that octamers harboring Sept9_i1 bind specifically and directly dynamic individual microtubules.

We then analyzed how the addition of Oct_9i1 or Oct_9i3, at 10 to 300 nM, affected parameters of microtubule dynamic instability (Fig. 7a, b). Both types of octamer were similarly and moderately affecting microtubule growth rate at intermediate concentrations. Strikingly, Oct_9i1 induced a strong and dose-dependent decrease of the microtubule shortening rate, while Oct_9i3 had no effect, even when used at the highest concentration. The catastrophe time frequency was not consistently altered by the addition of either types of octamer. Intriguingly, we observed that, solely in the presence of 200 nM or 300 nM of Oct_9i1, about 10% of depolymerizing microtubules had an unusual curved tubulin-containing structure at their plus-end that was decorated by septins (Fig. 7c and Supplementary Movie 3). These structures could be made of few protofilaments whose shortening is slowed down by the association with Oct_9i1.

**Fig. 7.**
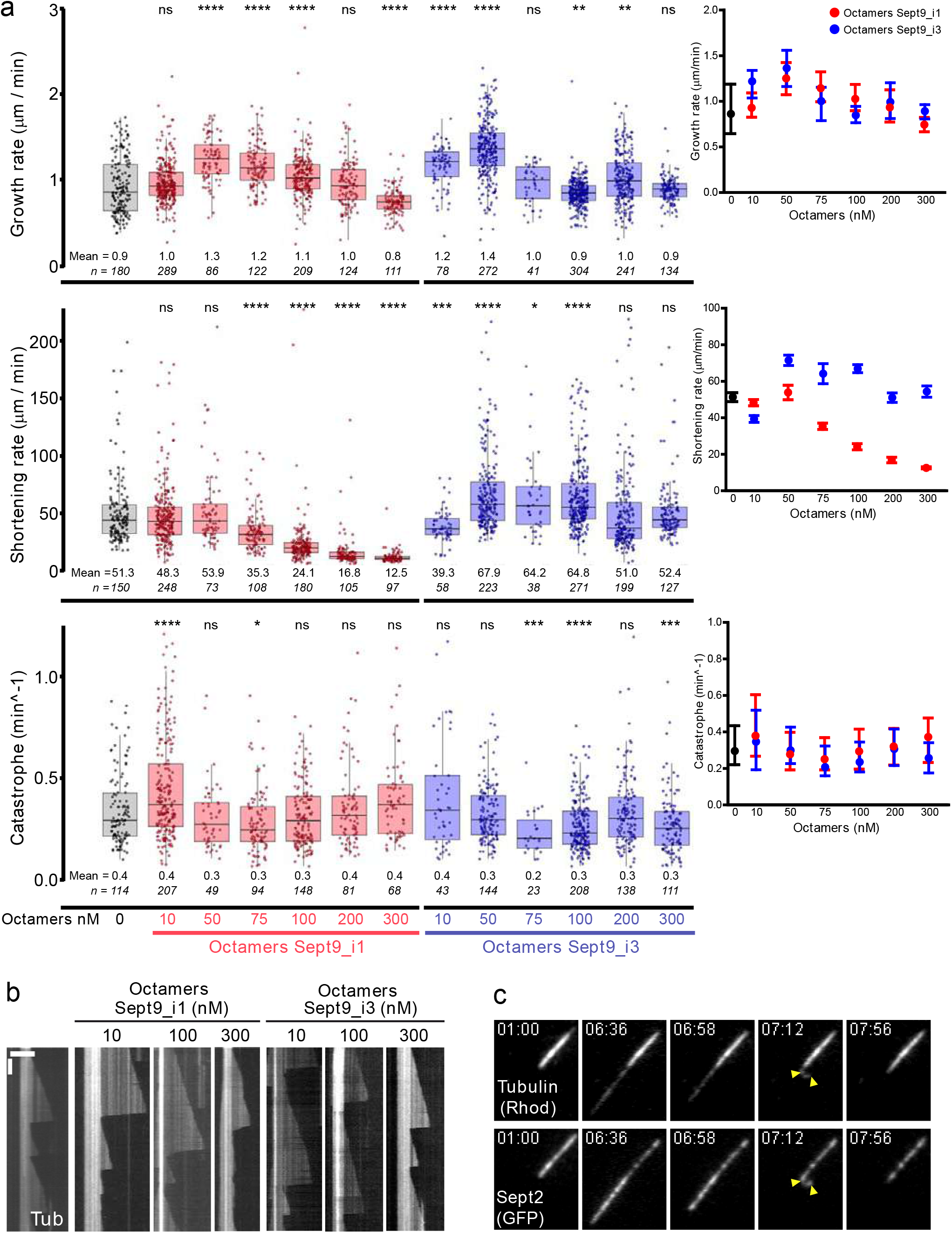
Recombinant septin octamers containing Sept9_i1 reduce the shortening rate of dynamic microtubules *in vitro*. **a** Quantitation of microtubule dynamic parameters (growth rate, shortening rate, and catastrophe time frequency) at increasing septin octamer concentrations. Results are presented as box and whiskers plots superposed with individual data points including outliers. The number of microtubules used (n) and mean value are indicated under each plot. (grey: no septin octamers, pink: octamers with Sept9_i1, light blue: octamers with Sept9_i3). Two-tailed t-test with Benjamini & Hochberg p-value correction with R, * p<0.05, ** p<0.01, *** p<0.001, **** p<0.0001. Right inset graphs show median values of each parameter in the absence of octamers (black dots) and direct comparison in presence of either octamers with Sept9_i1 (red dots) or octamers with Sept9_i3 (blue dots) and corresponding interquartile ranges. **b** Representative kymographs (rhodamine channel for tubulin) representing the effect of recombinant septin octamers on microtubule dynamics. Such kymographs were used to calculate the parameters values presented in **a** from individual microtubules. First kymograph on the left is in the absence of septin octamers. Horizontal white scale bar 5 μm; vertical white scale bar 50 sec. **c** Snapshots at indicated times (min:sec) of a microtubule and associated septin octamers harboring Sept9_i1 (200 nM, Supplementary Movie 3). Yellow arrow heads point at a curved structure on a depolymerizing microtubule (+) end. White scale bar 5 μm.

### Re-localization of octamers from stress fibers to microtubules alters stress fiber organization and function

Our observations indicate that the population Sept9_i1 octamers associate with both microtubules and actin fibers and can shuttle between them. This was for instance illustrated by the observation that the Sept9_i1-i5 chimera co-localized with microtubules only (Fig. 2f, h and Fig. 4b, c), but was able to co-localize with actin fibers upon nocodazole-induced depolymerization of microtubules (Fig. 4c). Thus, beyond its direct contribution to microtubule stability and dynamics (this study) and microtubule-linked processes, e.g. cell resistance to paclitaxel^52^, the relative distribution lof Sept9_i1 on microtubules vs actin fibers might indirectly affect actomyosin fibers distribution and cellular events dependent on their contractility. Previous studies showed that Sept9 associates with perinuclear actin fibers, especially ventral stress fibers whose organization depends on Sept9^40,53^ (Supplementary Fig. 5a). We similarly observed that in U2OS (Fig. 8a) and RPE1 cells (Supplementary Fig 5d), the knockdown of Sept9 strongly decreased the percentage of cells with sub-nuclear stress fibers. To test the hypothesis that septin octamer association with microtubules impacts actin fiber organization and function, we generated U2OS cell lines stably expressing Sept9_i1 variants (Supplementary Fig. 4).

**Fig. 8.**
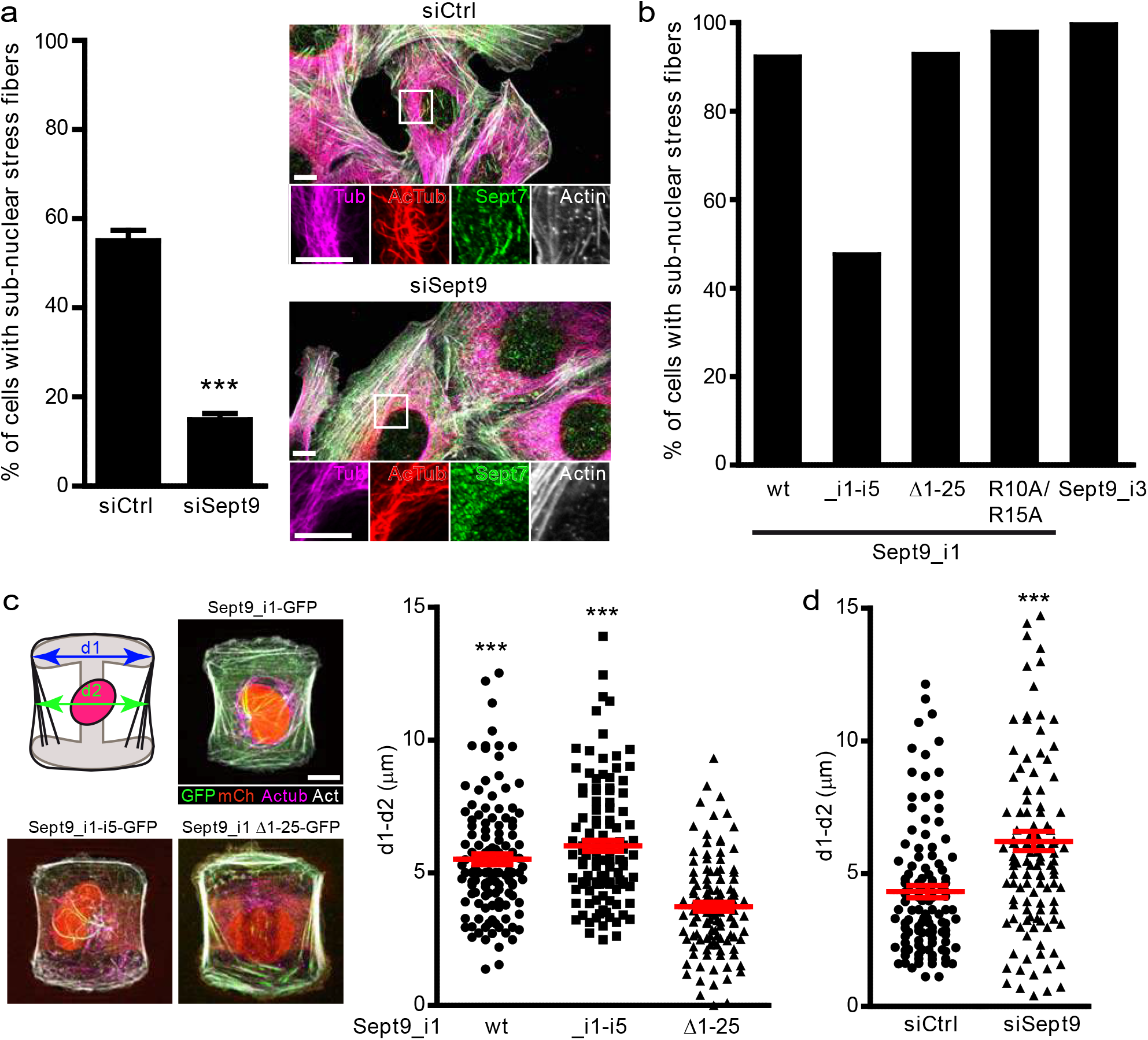
Impact on actin cytoskeleton and cell shape of septin octamer sequestration by microtubules via Sept9_i1 MBD. **a** Analysis of the presence of sub-nuclear actin fibers in U2OS transfected by either siCtrl or siSept9. Results are from three independent experiments on a total of 90 cells per condition (30 cells per experiments). Unpaired, two-tailed t-test with Welch’s correction, * p<0.05, ** p<0.01, *** p<0.0005, % of cells vs siCtrl. Right: Co-localization of Sept7 with sub-nuclear actin stress fibers. White scale bar 10 μm, insets: two-fold zooms of frames region. **b** Analysis of the presence of sub-nuclear actin fibers in U2OS derived cell lines stably expressing each one of the indicated C-terminally GFP tagged Sept9_i1 constructs and the histone mCherry-H2B fusion. Results are from a single determination on 100 cell images per cell lines. **c** Analysis of shape changes on derived U2OS cell lines described in **b** and adhering to medium size H micropatterns. d1: average maximum width of 35.170 μm observed from measurement on four empty H micropatterns; d2: observed width of single adherent cells at mid-maximal height of the H micropatterns. Representative images of merged fluorescence channels of the three U2OS derived cell lines selected for analyses. Note that as for the rest of the figure, these cell lines (U2OS.Sept9_i1-GFP.mCherry-H2B, U2OS.Sept9_i1-i5-GFP.mCherry-H2B and U2OS.Sept9_i1Δ1-25-GFP.mCherry-H2B) are indicated simply by the originally transfected Sept9-GFP construct names. White scale bar is 10 μm. Right: measurements of the difference between d1 and d2 in cells from selected lines showing a higher value when septins bind to microtubules, due to a reduction of the cell width at mid-maximal height. Results are from two independent experiments on a total of 120 cells (60 cells per experiment). Unpaired, two-tailed t-test with Welch’s correction, *** p<0.0005, d1-d2 vs Sept9_i1Δ1-25. Red lines represent mean values ± SEM. **d** Measurements of d1-d2, as described in c, in U2OS cells transfected either by either siCtrl or siSept9 as described in **c** showing that Sept9 KD reduced cell width at mid-maximal height. Results are from three independent experiments on a total of 120 cells (40 cells per experiment). Unpaired, two-tailed t-test with Welch’s correction, *** p<0.0005, d1-d2 vs siCtrl. Red lines represent mean values ± SEM.

Expression of Sept9_i1-i5, only associating with microtubules, was concomitant with the strong reduction in the percentage of cells having sub-nuclear stress fibers (Fig. 8b), while expression of Sept9 constructs with no or reduced microtubule association (Sept9_i1 Δ1-25, Sept9_i1 R10A/R15A and Sept9_i3) preserved sub-nuclear stress fibers (Fig.8b). Likewise, paclitaxel-induced re-localization of Sept9_i1 octamers from stress fibers to microtubule bundles in RPE1 cells induced the loss of sub-nuclear stress fibers (Fig. 5c), whereas paclitaxel-treatment of U2OS expressing Sept9_i3 as the major isoform did not result in any loss of sub-nuclear stress fibers (Fig. 5c). Thus, the sequestration of septin octamers on microtubules is dependent on the MBD of Sept9_i1 and induces a loss of sub-nuclear stress fibers. These results suggest that the precise relative distribution of septins on microtubules vs on actin fibers is critical for an intact actin cytoskeleton.

As a functional assay for testing the interplay between Sept9_i1-microtuble association and stress fiber function, we plated U2OS cells KD for Sept9 or expressing different Sept9_i1 variants on adhesive micropatterns. Indeed, when plated on adhesive micropatterns, cells form actin stress fibers across adhesives edges, and the convexity of the non-adhesive cell edges provides a readout for stress fiber tension^54^. Partial or total localization of septins on microtubules vs stress fibers in U2OS cells stably expressing Sept9_i1 or Sept9_i1-i5, respectively, promoted the concavity of non-adhesive cell edges, reflective of the relaxation of peripheral stress fibers compared to cells stably expressing Sept9_i1 Δ1-25, that is solely located on actin fibers (Fig. 8c). The knockdown of Sept9 in U2OS cells, leading to the loss of septin octamers, induced a similar detectable relaxation of peripheral stress fiber (Fig. 8d).

Altogether, these results suggest that the relative abundance of octamers vs hexamers and/or the relative proportion of septin octamers able to associate to microtubules contribute critically to the maintenance of the actomyosin organization and tension. Our study indicates that the propensity of octamers harboring Sept9_i1 to associate with microtubule depends on Sept9_i1 expression levels, the state of its N-terminal regulatory domain, and the extent of microtubule bundling.

## Discussion

Septins associate in a dynamic way with different major constituents of the cell, i.e. membranes, actin fibers and microtubules^6^. Septin octamers and hexamers co-associate to form septin filaments^7,8,10–13^, whose properties may depend in part on the relative abundance of incorporated octamers and hexamers. In addition, the expression profile of different septin isoforms, including those of Sept9, may direct both localization and function of septin octamers. It is thus critical to define precisely the molecular basis of the different subcellular localizations of septin complexes, in order to develop tools differentially altering their distribution. This will help us understand how the different septin pools respectively contribute to cell physiology and pathology.

Earlier studies have shown that Sept9_1 has a preferential affinity for microtubules in cells^24,32,40,42^ indicating that the first 25 aa residues defining the sequence unique to this long isoform was responsible for specific binding to microtubules. In the present study, we identify a conserved MAP-like MBD in Sept9_i1, including two distinct modules and key conserved residues whose mutation abrogates microtubule binding. We and others have identified imperfect repeats containing basic amino acid residues in the common long N-terminal sequence of Sept9_i1, _i2 and _i3^40,41^. These repeats were proposed to constitute the MBD of Sept9 via electrostatic interactions with the acidic C-termini of β-tubulin exposed at the surface of microtubules^41^, in line with the accepted mode of interaction of conventional MAPs at the time^55^. However, recent advances in the determination of the structure of MAP tandem repeats bound to microtubules by cryo-EM contradicted this model^46,47^. These studies revealed that a sequence in Tau and MAP4 repeats anchors MAPs to a binding pocket at the interface between consecutive tubulin heterodimers in protofilaments, independently of tubulin acidic C-termini. We actually found numerous similarities between the MAPs repeated sequences and the AIR9-like module of Sept9_i1 MBD, that are not present in the repeats shared with the other Sept9 long isoforms. Most importantly, our data show that, contrary to the Sept9_i1 MBD, the N-terminal region common to all long isoforms is not required for the association of Sept9_i1 harboring octamers with microtubules in cells. Deletion of this region actually favored septin association with microtubules at the expense of actin fibers. These findings led us to speculate that the common N-terminal domain is not directly involved in the interaction of Sept9_i1 with the microtubule lattice, but negatively regulates it. The imperfect repeats of the common N-terminal domain were predicted to form short β-sheets whereas the rest of the domain is disordered^40^. One of the imperfect repeats of the common N-terminal region is the locus of mutations detected in HNA patients^35–37^. To this date the mechanisms whereby Sept9 mutations contribute to this disease are not known. Remarkably, when we expressed Sept9_i1 harboring HNA-like mutations, septin octamers were no longer able to associate with microtubules, while still associating with actin fibers. This finding suggests that HNA-associated mutations alter the putative “Sept9_i1 MBD regulatory region” providing clues for the determination of the molecular mechanisms underlying HNA.

Importantly, we demonstrated that Sept9_i1 localizes septins on microtubules, only if it is included in polymerizable septin octamers. At endogenous expression levels, in the diverse cellular models we studied, we did not observe the presence of a significant pool of monomeric Sept9 (Fig. 1b), which suggests that microtubule-associated Sept9 is wholly incorporated in octamers in these cells. Conversely, the large excess of Sept9_i1 monomers in cells expressing the Sept9_i1 G interface mutant was not associated with microtubules (Fig. 3a, b). Thus, *in vitro* studies with isolated recombinant Sept9_i1 suggesting that Sept9_i1 can binds and bundle microtubules by itself ^41,56,57^ must be interpreted with caution.

In previous studies, recombinant Sept9_i3 or the common N-ter fragments were shown to associate with microtubules, assembled from purified tubulin^41,56^. The discrepancy is most likely linked to the fact that in these *in vitro* experiments, Sept9 was used outside of its natural octameric context, often at high concentrations (up to 10-30 μM in pelleting assays) and using paclitaxel or GMPCPP stabilized microtubules in very low salt buffers, which likely favored electrostatic interactions. Using recombinant octamers harboring Sept9_i1 and Sept9_i3^12^, we observed that only those harboring Sept9_i1 associated with dynamics microtubules at low nanomolar concentrations, *in vitro*. Our *in vitro* reconstitution experiments results are the first one to recapitulate the specific binding of Sept9_i1 octamers to microtubules observed in cells here and by others^40,52^. Of note, although recombinant Sept2-Sept6-Sept7-Sept7-Sept6-Sept2 hexamers were found to interact with microtubules *in vitro*^58^, this is not consistent with the absence of co-localization of septin hexamers with microtubules, when Sept9 was knocked down in cells^40^ (Fig 8a and Supplementary Fig. 1d).

Septins associate with a sub-population of bundled microtubules^41,42,59^. We confirm that Sept9_i1 co-localize with acetylated, bundled microtubules (much more selectively than structural MAPs, such as MAP4). As we also show that microtubule acetylation is not a requirement, Sept9_i1 binding appears rather to rely on microtubule bundling in cells. Intriguingly, microtubule bundling is not required *in vitro*. This paradox will require other studies, including structural ones, to further characterize the molecular bases of the association of septins with individual and bundled microtubules.

Our data show that octamers harboring Sept9_i1 slowed down microtubule depolymerization *in vitro* and stabilized microtubules in cells treated with nocodazole. It will be important to determine how these molecular events contribute to microtubule-dependent cellular processes such as spatiotemporal sorting of vesicles and organelles^48,57^, formation of protrusive structures^26–29,60,61^ or cell survival against cytotoxic doses of paclitaxel^52^.

Association of septins with microtubules is likely to be regulated by cell- and tissue-specific contexts. Indeed, septin association with microtubules is clearly correlated to the specific expression of the Sept9_i1 isoform and is sensitive to the bundling state of microtubules. Septin-microtubule interaction might also be modulated by the presence of competing MAPs or PTMs targeting the Sept9_i1 MBD and/or its putative regulatory domain. Importantly, we show that deficient Sept9_i1 binding to microtubules is associated with a diseased state, as two distinct HNA-like point mutations prevented Sept9 binding to microtubules. We are still far from understanding the underlying mechanism leading to HNA, which might be related to dysregulated microtubule function, but might be related also to an excessive association of septins with actin fibers and perturbation of actomyosin functions. Thus, our work not only establishes a new molecular basis for studying the physiological relevance of the association of septin with microtubules, but also open new avenues in the difficult search for the causative events leading to this rare neuropathy.

## Materials and methods

### Human Cell lines

U2OS (ATCC), SKBR-3 (ATCC), HeLa (a kind gift from Dr Patrice Dubreuil, CRCM, Marseille, France), RPE-1 and ARPE19 (kind gifts from Dr Michael Sebbagh, CRCM, Marseille) cells were maintained in Dulbecco’s modified Eagle’s medium (DMEM) (Gibco) supplemented with 4 mM GlutaMAX™, 10 % fetal bovine serum (FBS) and 1% sodium pyruvate (supplemented DMEM), except ARPE19 cells that were cultivated in DMEM/F12 (1:1 mixture of DMEM and Ham’s F12, Gibco) supplemented as for DMEM. All cell lines were maintained in the presence of 5% CO_2_ humidified atmosphere at 37°C. RPE-1 cells were also grown on coverslips coated with collagen (Corning, Cat#354236) until they reached confluency and ciliogenesis was induced by growing cells in DMEM containing only 0.5% of fetal bovine serum for 24 hours.

### Plasmids coding for fluorescent septin constructs and transfection

To drive expression of the constructs in mammalian cells, we used the immediate early enhancer and promoter of human cytomegalovirus (CMV promoter, 508 base pairs). Human Sept9_i1 cDNA was a gift from C. Montagna (Albert Einstein College of Medicine, USA). Human Sept9_i3 cDNA was a gift from W. Trimble (University of Toronto, Canada). A synthetic human Sept2 coding sequence (Eurofins Genomics, Germany) was generated using the codon usage of mouse Sept2 except for the five codons that differ between the two species, for which we used codons encoding the human residues. C-terminal green and red fluorescent protein fusions were generated using monomeric (V206K) superfolder GFP (msfGFP)^62–65^ and monomeric Apple (mApple)^64,66^, respectively. All constructs were generated with two-insert seamless cloning (In-Fusion HD Cloning Plus Kit from Takara Bio, Cat# 638910) using NheI/BamHI linearized plasmid backbones (Addgene plasmid #54759) and the oligonucleotide primer sequences (listed in Supplementary Table 1). Primers for seamless cloning were Cloning Oligo (<60 bp) or EXTREmer (>60 bp) synthesis and purification quality from Eurofins Genomics, Germany. Restriction enzymes were FastDigest enzymes from Thermo Scientific. All plasmids were verified by sequencing (Eurofins Genomics, Germany) after each cloning step, including the midipreps used for plasmid production. Constructs were transiently transfected by nucleofection in U2OS cells (Amaxa nucleofector program U2OS X-001, Lonza) as described previously^40^.

### siRNAs and transfection

The following 19-mer duplex siRNAs were purchased from LifeTechnologies: siRNA control (siCtrl) targeting the E.Coli β galactosidase (LacZ) (5’-GCGGCUGCCGGAAUUUACC-3’), Sept9 (5’-GGAUCUGAUUGAGGAUAAA-3’) targeting the 3’UTR of all human Sept9 mRNA variants, Sept7 (5’-CGACUACAUUGAUAGUAAA-3’) targeting the human mRNA coding region, αTAT1 (5’-CGCACCAACUGGCAAUUGA-3’), whose design was based on validated siRNA αTAT1#2 from Shida et al^67^. Transfection of siRNA was performed as described previously^40^ either with Lipofectamine RNAiMAX (Invitrogen, Cat# 13778075) or Amaxa cell line nucleofection kit V (Lonza, Cat#VCA-1003).

### Drug treatments

For some experiments, cells were treated for two hours with 2 μM paclitaxel (Sigma-Aldrich, Cat#T7191) or 10 μM nocodazole (Sigma-Aldrich, Cat#M1404) in DMSO or with DMSO alone (Sigma-Aldrich, Cat#D8418).

### Antibodies

Labeling of septins on Western blots (WB) and/or on immunocytochemistry coverslip (ICC) was achieved with the following antibodies: rabbit polyclonals against human Sept2 (Sigma-Aldrich, Cat#HPA018481, WB), against human Sept9 (Sigma-Aldrich, Cat#HPA042564 and HPA050627, WB and ICC; Proteintech, Cat#10769-1-AP, WB), against Septin 7 (IBL, cat#18991, WB and ICC), against Sept11 (Sigma-Aldrich, Cat#SAB2102111, WB), against human Sept5, against Sept8 (kind gifts from Barbara Zieger, University of Freiburg, Germany, WB), against Sept6 (a kind gift from Makoto Kinoshita (Nagoya University, Japan, WB), against Sept10 (Sigma, Cat#HPA047860, WB); rat monoclonals against human Sept9_i1 (clone 4D2A5, WB) or human Sept9_i3 (clone 1A6C2, WB), and against human Sept7 (clone 10A7, WB and ICC) were produced as described previously^40^ (WB). Other antibodies were against α-tubulin: mouse monoclonal (Sigma-Aldrich, DM1A Cat#05-829, WB and ICC), rat monoclonal (Invitrogen, YL1/2 Cat#MA1-80017, ICC), mouse monoclonal anti-acetylated K40 (Santa Cruz Biotechnology, 6-11B1 Cat#sc-23950, WB and ICC), against tubulin: mouse monoclonal anti-glutamylated tubulin (Adipogen, GT-335 Cat#AG-20B-0020-C100 and polyE Cat#AG-25B-0030-C050, WB). Labeling of actin was performed with phalloidin-Atto 390 (Sigma-Aldrich, Cat#50556, ICC) or phalloidin-TRITC (Sigma-Aldrich, Cat#P1951, ICC). Secondary antibodies conjugated with HRP were used for WB (Dako Agilent). For ICC application, and for WB application when indicated in figures, the following fluorophores conjugated to secondary antibodies were used: DYLIGHT405, AlexaFluor405, AlexaFluor488 and AlexaFluor594 (Jackson, Immunoresearch) and AlexaFluor 647 (Invitrogen).

### U2OS cell lines stably co-expressing mCherry-H2B and Sept9-GFP constructs

U2OS and HeLa cell lines stably expressing mCherry-H2B and a Sept9_X-GFP construct were generated by co-transfecting the plasmid pBabeD hygro mCherry-Histone H2B (a kind gift from Christophe Lachaud, CRCM, Marseille, France) with hygromycin B resistance and one of the plasmids encoding for different Sept9 isoforms and mutants-GFP constructs, with geneticin resistance. U2OS cells were nucleo-transfected (Amaxa nucleofector program U2OS X-001, Lonza) with Amaxa kit V. Three days after transfection, co-expressing cells were selected with hygromycin B (Invitrogen, Cat#10687-010) and geneticin (Gibco, Cat#10131-027) at 0.5 mg/mL each. GFP and mCherry fluorescence positive cells were sorted by flow cytometry. Sorted cell populations were cultured in supplemented DMEM in the presence of 0.5 mg/mL of antibiotics. After three passages, cell line stocks were stored in liquid nitrogen.

### Micropatterning of U2OS cells

6×10^3^ U2OS cells KD for Sept7 or Sept9 or U2OS cells stably co-expressing mCherry-H2B and Sept9-GFP constructs were seeded in each well formed by the assembly of a 35 mm collagen coated coverslip with H-shaped medium size adhesive micropatterns (Cytoo, Cat#10-008-00-18) and a magnetic four-well chamber (Cytoo, Cat#30-011) following manufacturer instructions. Coverslips were fixed with 4% formaldehyde and processed for ICC as previously described^40^. Distance measurements were performed using the Graphics line tool in Zen blue software as described in more details in Fig. 8c legend.

### SDS-PAGE and Western blotting

Total protein extraction from cells in culture by scraping the cells off in NP-40 lysis buffer, separation of proteins by SDS-PAGE using NuPAGE 4-12 % gradient Bis-Tris gels (Invitrogen, Cat#NP0322BOX) and MOPS SDS running buffer (Invitrogen, Cat#NP0001-02) and Western blotting of separated proteins on nitrocellulose were performed as described previously^40^.

### Separation of septin complexes by native gel electrophoresis and detection

Native cellular protein extraction was carried out as published by Sellin et al.^68^. Briefly, proteins from cells at 80% confluence in one 100 mm petri dish were extracted in 40 μl of native lysis buffer (80 mM PIPES pH 6.9, 2 mM MgCl2, 4 mM EGTA, 0.2% saponin and protease inhibitor cocktail (Roche, Cat#04693159001); the extract was incubated for 10 min on ice and centrifuged and the supernatant was supplemented with 0.45 M sodium chloride, incubated on ice, centrifuged, concentrated and exchanged twice with phosphate buffer (pH 7.5, 0.45 M NaCl, 1 mM EGTA and protease inhibitors) on an Amicon 30 kDa cut off concentrator (Merck, Cat# UFC 503096). Protein concentration was determined using a BCA protein assay (Thermo Scientific, Cat#23223 and 23224), glycerol was added at 1:1 (vol:vol), and extracts stored at −20°C until use. Native PAGE was performed on 4-16% NativePAGE Novex Bis-Tris polyacrylamide gels (Thermo Scientific, Cat#BN1002BOX) following instructions from the manufacturer. Briefly, native protein extracts were mixed with 4x sample buffer (Thermo Scientific, Cat#BN2003) and water (final sample buffer 1x, and about 150 mM NaCl). Eight μg of proteins were loaded per lane. Unstained native protein molecular weight standards were used (Thermo Scientific, Cat#LC0725) and stained with Coomassie blue during electrophoresis. Protein complexes were transferred overnight on a PVDF membrane under 20 V constant voltage. The membrane was partially distained with 25% MeOH and 10% acetic acid, positions of native protein molecular weight standards were marked, and the membrane was placed briefly in MeOH for complete distaining. The membrane was processed for Western blot detection following the protocol used for nitrocellulose membranes described previously^40^.

### Immunocytochemistry

Multiplex immunocytochemistry was carried out on cells cultured on collagen coated coverslips followed by 4% formaldehyde fixation as described previously^40^. Images were acquired on a Zeiss structured light ApoTome microscope equipped with a 63x/1.4 Plan Apochromat objective and an Axiocam MRc5 camera using AxioVision software or a Zeiss LSM880 META confocal microscope equipped with a 63x/1.46 Plan Apochromat objective and a GaAsP detector using Zen software.

### Co-localization of septins with microtubules and with stress fibers

Individual cells were examined by ICC for co-localization of septins with total α-tubulin, acetylated K40 α-tubulin and F-actin, from top to bottom. Cells were counted as positive for co-localization of septins with microtubules or with F-actin when septin fluorescence signal was co-aligned with microtubules and/or acetylated microtubules or with ventral stress fibers, respectively. The percentage of cells positive for both septin-microtubule and septin-stress fibers co-localization, only positive for septin-stress fibers co-localization and only positive for septin-microtubule co-localization were designated in graphs as Microtubule+Actin, Actin, and Microtubule, respectively. The extent of the co-localizations in each cell was not considered.

### Quantification of acetylated microtubules in cells

The fluorescence surface corresponding to acetylated microtubules in individual cells was determined using the ImageJ 1.48v software^69^: the fluorescence image of acetylated microtubules was converted in 8-bits, adjusted manually with the threshold function for black and white levels to maximally remove black pixels unrelated to acetylated microtubules, the actin image was synchronized with the acetylated microtubule image (Synchronize Windows tool) and used to delimit the ROI corresponding to the entire cell surface with the freehand selection tool, and finally the image was analyse using the measure function to obtain the percentage of surface area represented by the remaining black pixels corresponding to acetylated microtubule.

### Production and purification of recombinant human septin complexes

Human septin hexameric and octameric complexes with either Sept9_i1 or Sept9_i3 were produced, purified and analyzed as detailed in Iv et al^12^. No difference in apparent MW was observed between recombinant septin octamers or hexamers and those extracted from cells after native gel separation and WB (Supplementary Fig. 1b, c).

### Septin/microtubule reconstitution and TIRF imaging

GMPCPP-stabilized microtubule seeds serving as a nucleation site for dynamic microtubules were prepared using an established double-cycle method^70^. Briefly, a ~22 μM mixture of tubulin (~11 μM tubulin dimers) in MRB80 (80 mM PIPES pH 6.8, 4 mM MgCl2, 1 mM EGTA), composed of 75% unmodified tubulin dimers (Cytoskeleton, Inc., Ct#T240), 15% rhodamine-labeled tubulin dimers (Cytoskeleton, Inc., Cat#TL590M) and 10% biotin-modified tubulin dimers (Cytoskeleton, Inc., Cat#T333P), was spun down using an Airfuge^®^ Air-driven ultracentrifuge (Beckman Coulter Inc., Brea, California, USA) for 5 minutes at 30 psi with a cold rotor. Then, the mixture was complemented with 1 mM GMPCPP from a 10 mM stock solution, thus diluting the tubulin to 20 μM. This mixture was incubated at 37°C for 30 minutes to polymerize tubulin, and immediately airfuged for 5 minutes at 30 psi with the rotor at room temperature to pellet GMPCPP-stabilized microtubules. Afterwards, the supernatant was discarded and the pellet was resuspended in warm MRB80 to a final tubulin concentration of 20 μM, considering an 80% recovery of the tubulin. The mixture was incubated on ice for 20 minutes to depolymerize the microtubules. Subsequently, it was complemented with 1 mM GMPCPP, incubated at 37°C for 30 minutes to repolymerize the microtubules, and airfuged for 5 minutes at 30 psi with a warm rotor. The pellet, containing GMPCPP-stabilized microtubule seeds, was resuspended in warm MRB80 supplemented with 10% glycerol, snap-frozen, and kept at −80°C until use.

Nr. 1 Menzel coverslips (Thermo Fisher scientific, product number 11961988) and glass slides (Thermo Fisher scientific, product number 11879022) were cleaned in base piranha solution (5% hydrogen peroxide, 5% ammonium hydroxide) at 70°C for 10 minutes, washed extensively by rinsing with Milli-Q water and stored in Milli-Q water for up to 5 days. Just before use, a coverslip and a glass slide were blow dried with a stream of N2 gas. Flow channels were prepared by placing parallel 2×20 mm parafilm strips spaced by 2-3 mm between the glass slide and the coverslip. The parafilm was melted by placing the chambers on a hotplate at 120°C. After cooling down, the chambers were passivated by incubation with 0.2 mg/mL Poly(L-lysine)-graft-biotinylated PEG (SuSoS, product bumber CHF560.00) in MRB80, 0.2 mg/mL neutravidin in MRB80, 0.5mg/mL κ-casein in MRB80, and 1 wt% Pluronic F-127 in MRB80, in that order and without intervening washing steps. The channels were washed with three channel volumes (~30 μL) of MRB80 at the end of the passivation process. Then the channels were incubated for 10 minutes with an aliquot of GMPCPP-stabilized seeds that was quickly thawed at 37°C, allowing the microtubule seeds to bind to the surface via biotin-neutravidin interactions. Microtubule polymerization was induced by immediately flushing into the channel 20 μM tubulin (3% rhodamine-labeled tubulin) in a 5:1 volume ratio of MRB80:septin buffer complemented with 0.5 mg/mL κ-casein to prevent unspecific interactions, 0.1% methylcellulose as a crowding agent, 1 mM ATP, 1 mM GTP and an oxygen scavenging system composed of 50 mM glucose, 200 μg/ml catalase, 400 μg/ml glucose-oxidase, and 4 mM DTT. A mixture of 90% unlabelled and 10% msGFP-labeled septin hetero-hexamers or hetero-octamers (containing either SEPT9_i1 or Sept9_i3) were added to the previous mix at different concentrations in the range of 10 to 300 nM.

The samples were immediately imaged using a Nikon Ti2-E microscope complemented with a Gataca iLAS2 azimuthal TIRF illumination system heated to 30°C using an Okolab incubator system. The sample was illuminated with 488-nm and 561-nm lasers (Gataca laser combiner iLAS2) to visualize the septin and the tubulin signals, respectively. To achieve fast imaging, the fluorescence signal was split with a Cairn Research Optosplit II ByPass containing a Chroma ZT 543 rdc dichroic mirror and filtered with either a 525/50 or a 600/50 chroma bandpass filter. The images were recorded with a Andor iXon Ultra 897 EM-CCD camera using an exposure time of 75 ms, for 10-20 minutes with a frame rate of 1 frame/second.

Microtubule dynamics were analysed by kymograph analysis^71^. Kymographs were built with the reslice tool in FIJI^69^ on a manually drawn line that went from the beginning of the microtubule seed to the tip of the microtubule in its maximum length. In the kymographs, we can observe the positions of the microtubule tips over time. The analysis was only done on the plus ends of the microtubules, which can be distinguished from the minus ends via the longer final length and higher growth velocity of the former. Growth and shortening rates were obtained as the slopes of manually fitted straight lines on the growing or shortening contour phases, determined by manual inspection, of the microtubule plus ends. The catastrophe rate was calculated as the inverse of the time that a microtubule spends growing.

### Statistical analysis

Unless indicated otherwise, statistical analysis was performed using GraphPad Prism software and unpaired two-tailed t-test with Welch correction for cell related data or using the function compare_means from the package ggpubr from R and two-tailed t-test with Benjamini & Hochberg p-value correction (https://rpkgs.datanovia.com/ggpubr/index.html) for the *in vitro* data related to microtubule dynamics.

## Acknowledgements

This research received funding from the Agence Nationale pour la Recherche (ANR grant ANR-17-CE13-0014; SEPTIMORF) (to M.M. and P. V-P.). We thank Jeffrey den Haan (TU Delft, The Netherlands) for help with protein purification. This work was further financially supported by the Netherlands Organization for Scientific Research (NWO/OCW) through the ‘BaSyC-Building a Synthetic Cell’ Gravitation grant (024.003.019) (to G.K.).

## Authors contributions

J. Fialová, F. Iv, A. Llewellyn, M. Belhabib, M. Gomes, Y. Liu, K. Asano, D. Salaün: investigation; M. Kuzmić, G. Castro Linares investigation, writing – review & editing; T. Tachibana, supervision, resources, funding acquisition; G. H. Koenderink, A. Badache, conceptualization, funding acquisition, supervision, writing – review & editing; M. Mavrakis, investigation, conceptualization, methodology, funding acquisition, supervision, writing – review & editing; P. Verdier-Pinard, investigation, conceptualization, methodology, funding acquisition, supervision, writing – original draft, writing – review & editing.

## Notes

### Competing Interest Statement

The authors have declared no competing interest.

