## Supplemental legends-Table1-Figs for "Septin-microtubule association requires a MAP-like motif unique to Sept9 isoform 1 embedded into septin octamers"

**Supplementary Figure legends**

**Supplementary Fig. 1. Separation of septin complexes on native blue gels and detection by Western blot. a** Relation between migration distances of native septin complexes from the top of the 4-16% acrylamide gradient gels and those of native MW markers; each data point represents the average of two or three independent determinations. Right: table presenting observed apparent MW and theoretical MW of the different complexes. **b** Recombinant septin hexamers and octamers were run on gradient native gels as in **a** and were detected by Coomassie blue staining; the asterisk indicates the migration position of putative septin monomers. **c** Native protein extracts from Hela cells were run on native gels and blotted on a PVDF membrane; septin hexamers and octamers were detected by WB. **d** U2OS cells were KD for Sept7 or Sept9 and native extracts were analyzed as in **c**.

**Supplementary Fig. 2. Protein sequence conservation of the Sept9_i1 MBD. a** Alignment of Sept9_i1 sequences grouped by phylogenetically closely related organisms (birds and reptiles highlighted in green) and mammals (quadrupeds in blue, rodents in purple, and primates in turquoise). Amino acid residues are highlighted based on Clustal X color coding. A conservation score histogram for each position is presented below the alignments of sequences; under each column is indicated the numerical index reflecting the conservation of physico-chemical properties of amino acid residues (asterisk : 100% identical residues, plus sign: alternative residues with conserved properties). Columns exhibit a color shading coding from the maximal conservation index (11=asterisk) in pure yellow to the minimal conservation index (2) in dark brown. Sept9_i1 1-25 sequences were recovered using the PSI-BLAST tool and the human sequence as the seed, imported in Jalview, aligned with Mafft (redundant and truncated sequences were eliminated), and grouped by phylogenic proximity by BLOSUM52; the two highly conserved arginine residues R10 and R15 mutated in this study are indicated at the top of alignments. **b** Consensus logo of the Sept9_i1 MBD was generated using Jalview.

**Supplementary Fig. 3. Cellular localization of Sept9 by immunocytochemistry. a** Fluorescent detection of Sept9 and tubulin on Western blots of total protein extracts from U2OS cells transiently transfected by a control siRNA (siCtrl) or co-transfected with an siRNA against the 3'UTR region of Sept9 mRNA (siSept9) and with respective constructs described in Fig 2b. Numbers at the bottom of each lane indicate the levels of expression of Sept9 constructs relative to those of endogenously expressed Sept9 in first lane as quantified on this WB. **b** Percentages of U2OS cells transfected with the indicated Sept9-GFP constructs displaying co-localization of Sept9 with microtubules and actin fibers or only actin fibers during cytokinesis. Percentages were determined from three independent experiments based on a total of 90 cells (30 cells per experiment). Unpaired, two-tailed t-test with Welch's correction, * $p<0.05$, microtubule and actin localization vs Sept9_i1 wt. **c** U2OS cells transfected with the indicated Sept9-GFP constructs in cytokinesis. Insets are two-fold zoomed regions framed by a white square in the corresponding original images. **d** Percentages of U2OS cells transfected with the indicated Sept9-GFP constructs displaying co-localization of Sept9 with microtubules and actin fibers or only actin fibers during cytokinesis. Percentages were determined from three independent experiments based on a total of 90 cells (30 cells per experiment). Unpaired, two-tailed t-test with Welch's correction, ** $p< 0.01$, microtubule and actin localization vs Sept9_i5. **e** U2OS cells expressing Sept9_i5-GFP. Insets are 2.5-fold zoomed regions framed by a white square in the corresponding original images. **f** Comparison of percentages of cells displaying co-localization of Sept9 with microtubules and actin fibers or only actin fibers during cytokinesis in RPE1 and ARPE19 cell lines. Percentages were determined from three independent experiments based on a total of 90 cells (30 cells per experiment). Unpaired, two-tailed t-test with Welch's correction. **g** U2OS cells transfected with the indicated Sept9-GFP constructs and treated either by vehicle (DMSO) or 10 μM nocodazole for two hours.

**Supplementary Fig. 4. Characterization of U2OS cell lines stably co-expressing Sept9-GFP and mCherry-H2B constructs. a** Expression of Sept9-GFP constructs (Sept9_X-GFP) and total endogenous Sept9 (top Western blot) and expression of Sept9_i3-GFP constructs and endogenous

Sept9_i3 (bottom Western blot) in U2OS cell lines stably co-expressing mCherry-H2B and Sept9-GFP constructs (U2OS.Sept9_X-GFP.mCherry-H2B abbreviated here as U2OS.Sept9_X-GFP) and in the non-transfected parental U2OS cell line. **b** Percentages of cells displaying co-localization of Sept9_X-GFP with microtubules and actin fibers in U2OS.mCherry-H2B.Sept9_X-GFP cell lines. Results are from one determination in 30 cells from each cell line. **c** Images of Sept9_X-GFP constructs localization in U2OS.Sept9_X-GFP.mCherry-H2B cell lines, mCherry-H2B and acetylated tubulin fluorescence signals are shown in false blue and red colors, respectively.

**Supplementary Fig. 5. Phenotyping cells based on the presence of sub-nuclear actin fibers. a** Top and bottom view of actin fibers (phalloidin-Atto 390), Sept9_i1 (GFP), and nucleus (mCherry-H2B) in a U2OS.Sept9_i1-GFP.mCherry-H2B cell. White scale bar 10 µm. **b** Effects of KD of Sept7 or Sept9 in RPE1 cells on the relative expression of Sept7 and Sept9 and **c** on the presence of hexamers and octamers. **d** presence of sub-nuclear actin fibers in RPE1 cells KD for Sept9, i.e. in the absence of octamers. In **b**, Western blot signals of Sept7 or Sept9 expression were first normalized to the respective Western blot signals of α-tubulin and and then further normalized to the ratio obtained from cells transfected with Ctrl siRNA; results are from three independent experiments. Paired, two-tailed t-test with Welch's correction, ** $p< 0.01$, *** $p<0.0005$. In **c**, Western blots of native extracts resolved on native 4-16% acrylamide native gels. In **d**, results are from three independent experiments on a total of 90 cells (30 cells per experiment). Unpaired, two-tailed t-test with Welch's correction, ** $p< 0.01$, % of cells vs siCtrl.

**Supplementary Table 1. Oligonucleotide primer sequences used for the generation of the plasmids used in this study.** Oligonucleotides marked with an asterisk are shared among several constructs, but are listed only once.

**Supplementary Movie 1. Dynamics of septin octamers harboring Sept9_i1 (Oct_9i1) binding with microtubules *in vitro*.** Series of videos corresponding to each of the 10 to 300 nM of Oct_9i1 that were used. White scale bar 5 µm.

**Supplementary Movie 2. Dynamics of septin octamers harboring Sept9_i3 (Oct_9i3) binding with microtubules *in vitro*.** Series of videos corresponding to each of the 10 to 300 nM of Oct_9i3 that were used. White scale bar 5 µm.

**Supplementary Movie 3. Depolymerizing microtubules (examples 1 and 2) in presence of 200 nM of septin octamers harboring Sept9_i1 (Oct_9i1) *in vitro*.** Series of videos corresponding to each example. White scale bar 5 µm.

| promoter | in-house plasmid name | plasmid | primer name | primer sequence (5' to 3') | primer length |
|---|---|---|---|---|---|
| CMV | | Sept9_i1-msfGFP | NheISept9_i1FOR | CGTCAGATCCGCTAGCATGAAGAAGTCTTACTCAGGAGGCAC | 42 |
| | | Sept9_i3-msfGFP | NheISept9_i3FOR | CGTCAGATCCGCTAGCATGGAGAGGGACCGGATCTC | 36 |
| | | | Sept9sfGFPREV | TTGGACACCATCTCTGGGGCTTCTGGCTCC | 30 |
| | | | Sept9sfGFPFOR | AGAGATGGTGTCCAAGGGCGAGGAGC | 26 |
| | | | BamHIsfGFPREV | TAGATCCGGTGGATCCTTACTTGTACAGCTCATCCATGCC | 40 |
| CMV | | Sept2-mApple | NheIhSept2FOR | CGTCAGATCCGCTAGCATGTCTAAGCAACAACCAACTCAGT | 41 |
| | | | hSept2mAppleREV | CTCGCCCTTGCTCACCACATGGTGGCCGAGAGC | 33 |
| | | | mAppleFOR | GTGAGCAAGGGCGAGGAG | 18 |
| | | | BamHImAppleREV | TAGATCCGGTGGATCCTCACTTGTACAGCTCGTCCATGCC | 40 |
| CMV | pl-14 | Sept2 F20D V27D-mApple | NdeICMVpromoterFOR | ATCAAGTGTATCATATGCCAAGTACGCCCCCTATTGACGTCAATGACG | 48 |
| | | | Sept2FDREV | GATTGGGAAGATTTGCATCTCCAACATAGCCAGGAGTTTCTGG | 43 |
| | | | Sept2FDFOR | CAAATCTTCCCAATCAAGATCACCGAAAATCAGTGAAGAAGGGGTTCG | 48 |
| CMV | pl-6 | Sept9_i1-I281D M288D-msfGFP (Sept9_i1 NCmut2) | Sept9IDREV | TGCTCCAGGATGGAGTCATCCCCCACGTAGCCGAAGTCC | 39 |
| | | | Sept9MDFOR | CTCCATCCTGGAGCAGGATCGCCGGAAGGCCATGAAGC | 38 |
| CMV | α0h-4 | Sept9_i1 R289A R290A K291A K294A-msfGFP (Sept9_i1 NCmut1) | a0h_Sept9_REV | CGCCGCGGCCATCTGCTCCAGGATGGAGTCAATCC | 35 |
| | | | a0h_Sept9_FOR | CAGATGGCCGCGGCGGCCATGGCGCAGGGCTTCGAGTTCAACATCATG | 48 |
| CMV | pl-12 | Sept9_i1 W520A H530D-msfGFP (Sept9_i1 Gmut) | Sept9i1W520AREV | AACTTCGATGGTACCCGCCTTGGTCTTCCTCCCAAGGATCC | 41 |
| | | | Sept9i1H530DFOR | GGTACCATCGAAGTTGAAAACACCACAGATTGTGAGTTTGCCTACCTGCGG | 51 |
| CMV | is-6 | Sept9δN-msfGFP (Sept9_i1 Δ1-25) | NheISept9dNFOR | CGTCAGATCCGCTAGCATGGCCTTGAAAAGATCTTTTGAGGTCG | 44 |
| CMV | is-7 | Sept9_i5-msfGFP | NheISept9i5FOR | CGTCAGATCCGCTAGCATGGCCGACACCCCCAG | 33 |
| CMV | is-9 | Sept9_i1-i5-msfGFP | NheI9i1_Sept9i5FOR | CGTCAGATCCGCTAGCATGAAGAAGTCTTACTCAGGAGGCACGCGGACCTCCAGTGGCCGGCTCCGGAG GCTTGGTGACTCCAGTGGCCCAGCCGACACCCCCAGAGATG | 110 |
| CMV | mt-1 | Sept9_i1 Δ10-25-msfGFP | NheISept9i1D10FOR | CGTCAGATCCGCTAGCATGAAGAAGTCTTACTCAGGAGGCACGGCCTTGAAAAGATCTTTTGAGGTCGA GGAG | 73 |
| CMV | mt-2 | Sept9_i1 R10A R15A-msfGFP | NheISept9i1R10R15FOR | CGTCAGATCCGCTAGCATGAAGAAGTCTTACTCAGGAGGCACGGCCACCTCCAGTGGCGCCCTCCGGAG GCTTGGTGACTCC | 82 |
| CMV | mt-5 | Sept9_i1 S12A S13A-msfGFP | NheISept9i1S12S13FOR | CGTCAGATCCGCTAGCATGAAGAAGTCTTACTCAGGAGGCACGCGGACCGCCGCCGGCCGGCTCCGGAG G | 70 |
| CMV | mt-6 | Sept9_i1 S22A S23A-msfGFP | NheISept9i1S22S23FOR | CGTCAGATCCGCTAGCATGAAGAAGTCTTACTCAGGAGGCACGCGGACCTCCAGTGGCCGGCTCCGGAG GCTTGGTGACGCCGCCGGCCCAGCCTTGAAAAGATCTTTTGAGG | 113 |
| CMV | mt-20 | Sept9_i1 ΔN7-msfGFP (Sept9_i1 Δ1-7) | NheISept9i1DN7FOR | CGTCAGATCCGCTAGCATGGGCACGCGGACCTCCAG | 36 |
| CMV | Hna-1 | Sept9_i1 R106W-msfGFP | Sept9RWREV | CAGTGCGCCAGGACACCGGCTCGGCC | 26 |
| | | | Sept9RWFOR | TGTCCTGGCGCACTGAGCTGTCCATTGAC | 29 |
| CMV | Hna-2 | Sept9_i1 S111F-msfGFP | Sept9SFREV | GATGTCAATGAACAGCTCAGTGCGCCGG | 28 |
| | | | Sept9SFFOR | CTGTTCATTGACATCTCGTCCAAGCAGGTG | 30 |

**Supplementary Table 1. Oligonucleotide primer sequences used for the generation of the plasmids used in this study.**

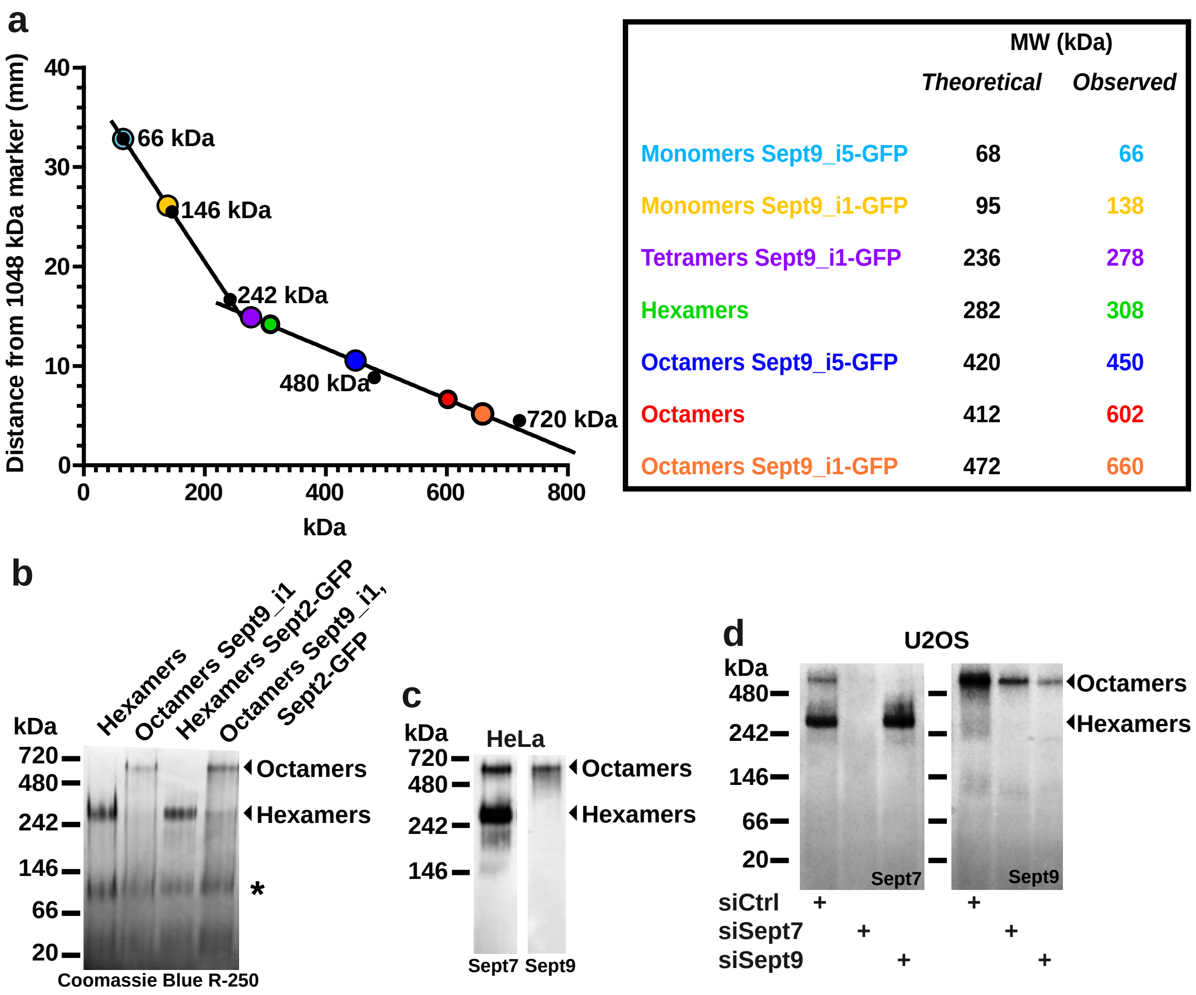


Kuzmic et al. Supplementary Figure 1

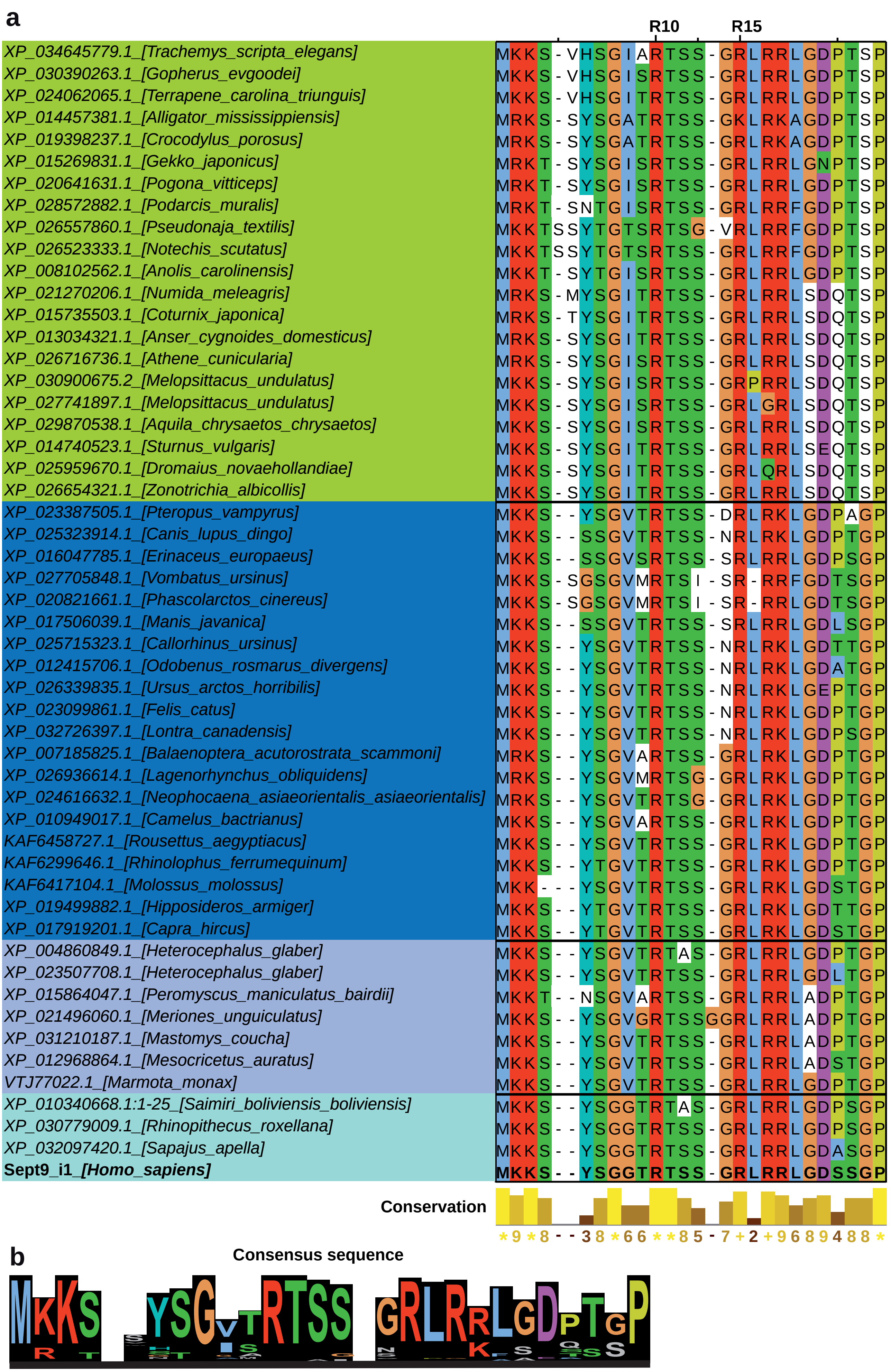


Kuzmic et al. Supplementary Figure 2

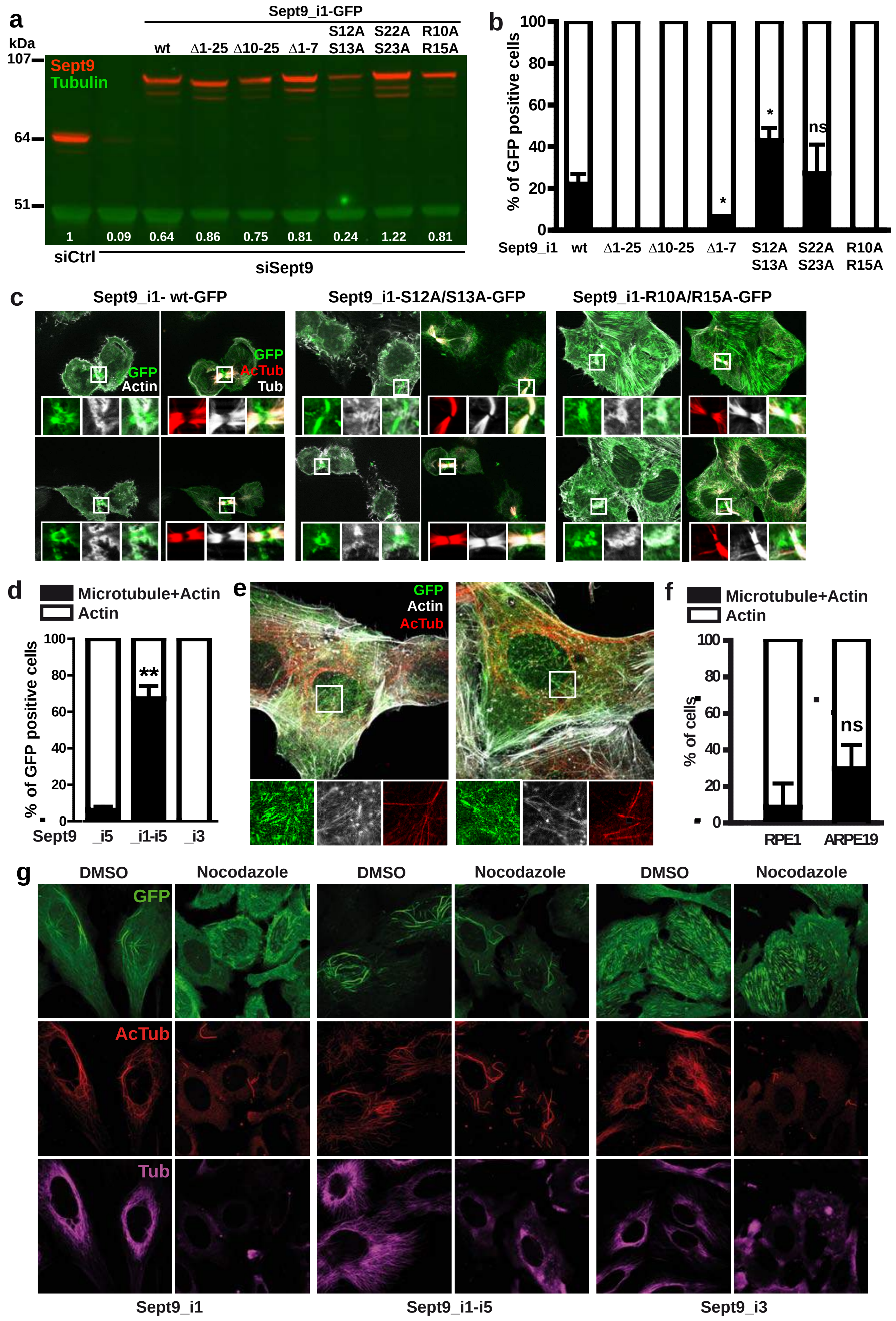


Kuzmic et al. Supplementary Figure 3

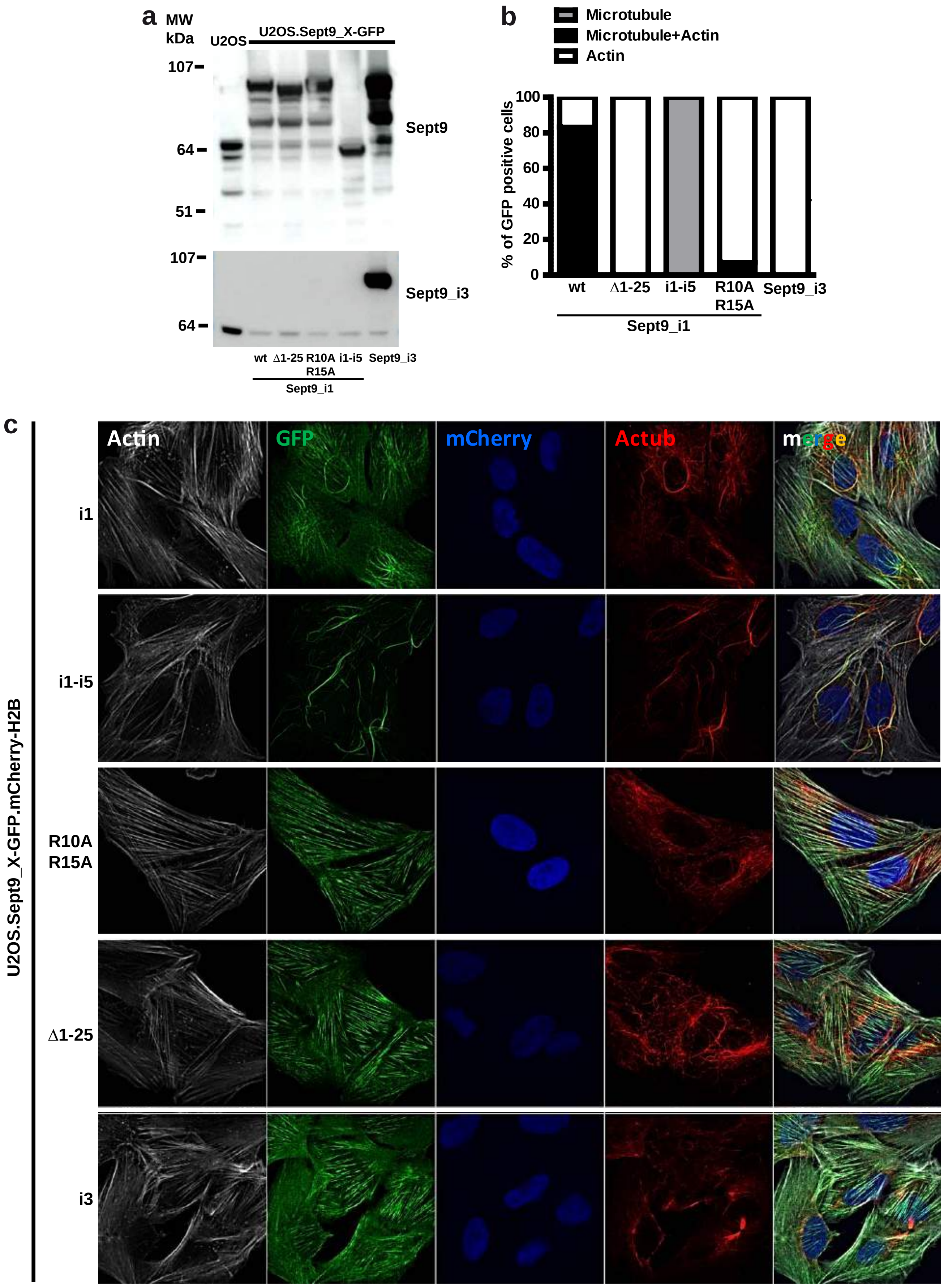


Kuzmic et al. Supplementary Figure 4

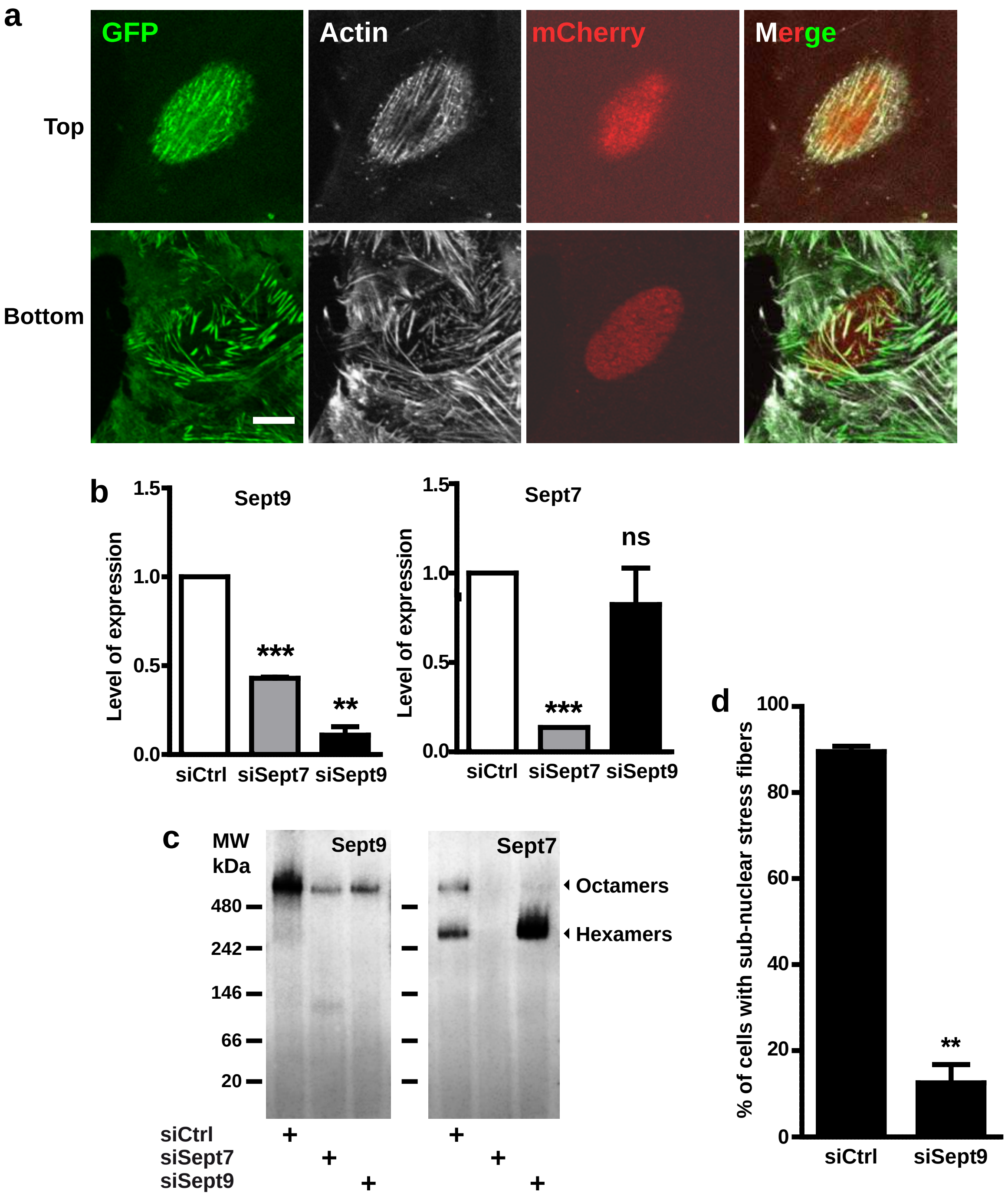


**Kuzmic et al. Supplementary Figure 5**
